# Myeloid *Zfhx3* Deficiency Protects Against Hypercapnia-induced Suppression of Host Defense Against Influenza A Virus

**DOI:** 10.1101/2023.02.28.530480

**Authors:** S. Marina Casalino-Matsuda, Fei Chen, Francisco J. Gonzalez-Gonzalez, Hiroaki Matsuda, Aisha Nair, Hiam Abdala-Valencia, G. R. Scott Budinger, Jin-Tang Dong, Greg J. Beitel, Peter H. S. Sporn

## Abstract

Hypercapnia, elevation of the partial pressure of CO_2_ in blood and tissues, is a risk factor for mortality in patients with severe acute and chronic lung diseases. We previously showed that hypercapnia inhibits multiple macrophage and neutrophil antimicrobial functions, and that elevated CO_2_ increases the mortality of bacterial and viral pneumonia in mice. Here, we show that normoxic hypercapnia downregulates innate immune and antiviral gene programs in alveolar macrophages (AMØs). We also show that zinc finger homeobox 3 (Zfhx3), mammalian ortholog of zfh2, which mediates hypercapnic immune suppression in *Drosophila*, is expressed in mouse and human MØs. Deletion of *Zfhx3* in the myeloid lineage blocked the suppressive effect of hypercapnia on immune gene expression in AMØs and decreased viral replication, inflammatory lung injury and mortality in hypercapnic mice infected with influenza A virus. Our results establish Zfhx3 as the first known mammalian mediator of CO_2_ effects on immune gene expression and lay the basis for future studies to identify therapeutic targets to interrupt hypercapnic immunosuppression in patients with advanced lung diseases.

**Graphical abstract:** 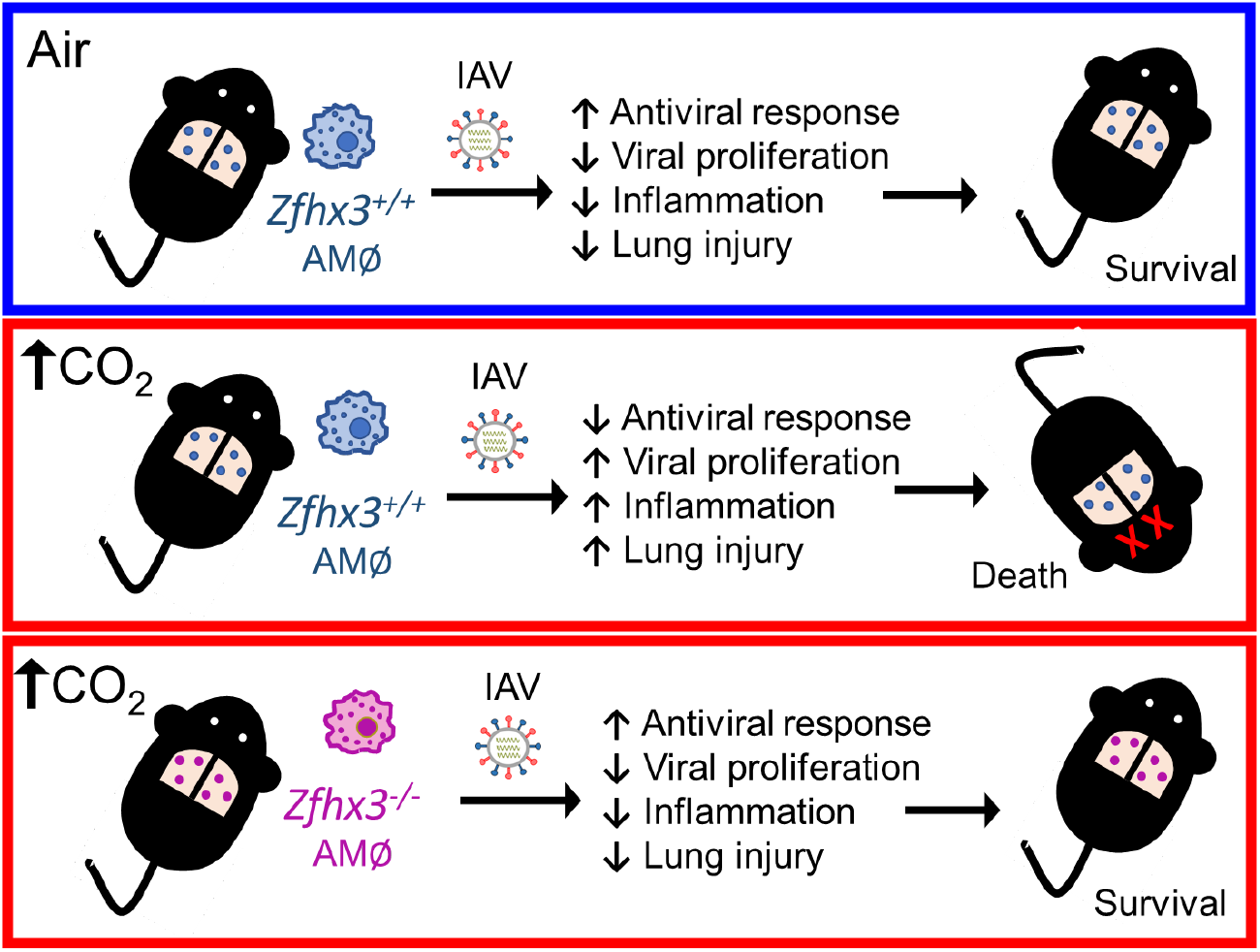

## Introduction

Hypercapnia, elevation of PCO_2_ in blood and tissue, commonly occurs in advanced chronic obstructive pulmonary disease (COPD) and in acute respiratory failure. Patients with severe COPD frequently develop bacterial and viral lung infections ^1^, including influenza ^2-4^, and hypercapnia is a risk factor for mortality in such individuals ^5-9^. Hypercapnia is also an independent risk factor for mortality in community-acquired pneumonia ^10^, adenoviral lung infections ^11^ and cystic fibrosis ^12^. Previously, we reported that hypercapnia suppresses transcription of multiple NF-kB-regulated innate immune genes required for host defense and inhibits phagocytosis in human, mouse, and *Drosophila* cells ^13-15^. We also showed that hypercapnia inhibits autophagy-mediated bacterial killing by macrophages (MØs) ^16^. Moreover, we found that hypercapnia increases mortality due to bacterial infections in both mice ^17^ and *Drosophila* ^13^. Recently we showed that normoxic hypercapnia inhibits antiviral gene and protein expression and increases viral replication, lung injury and mortality in mice infected with influenza A virus (IAV) ^18^. The suppressive effect of elevated CO_2_ on antiviral gene and protein expression and the hypercapnia-induced increase in IAV replication were particularly striking in lung MØs and were mediated by CO_2_-induced activation of Akt1 ^18^.

The similarity of hypercapnia’s effects in the *Drosophila* and mammalian systems suggested that elevated CO_2_ inhibits innate immune gene expression by conserved pathway(s). To identify putative molecular mediator(s) of such pathway(s), we conducted a genome-wide RNAi screen for genes whose expression was required for hypercapnic suppression of an antimicrobial peptide in cultured *Drosophila* cells. The screen identified multiple candidate CO_2_ mediators, the most potent of which was *zfh2*, a zinc finger homeobox transcription factor not previously known to have immunoregulatory function ^19^. Strikingly, mutant *Drosophila* deficient in *zfh2* were protected against the CO_2_-induced increase in mortality from bacterial infection ^19^. These results identified Zfh2 as the first known mediator of hypercapnic immune suppression in vivo. Like *Drosophila* Zfh2, its mammalian ortholog, zinc finger homeobox 3 Zfhx3^1^, also known as AT-binding transcription factor 2 (Atbf1), is a very large zinc-finger homeodomain transcription factor. Two isoforms, Zfhx3-A (MW 404 kD) and Zfhx3-B (MW 306 kD) are expressed as result of alternative promoter usage combined with alternative splicing^20^. Zfhx3 regulates neuronal differentiation^21^, functions as a tumor suppressor^22^ and sequence variants of the gene are associated with atrial fibrillation^23^. In the mouse, homozygous germline deletion of *Zfhx3* is embryonic lethal, and heterozygosity results in partial fetal loss and high neonatal and pre-weaning mortality ^24^. Lung MØs are critical for resistance to influenza virus infection ^25-28^. This, along with our findings that hypercapnia downregulates MØ antiviral gene and protein expression ^18^ and that Zfh2 deficiency abrogates hypercapnic suppression of resistance to bacterial infection in *Drosophila* ^*19*^, led us to hypothesize that Zfhx3 might mediate the effects of elevated CO_2_ on lung MØs and host defense against IAV. Here we show that Zfhx3 is highly expressed in mouse and human MØs. We thus bred a mouse selectively lacking *Zfhx3* in the myeloid lineage, which in the unstressed state grows normally without apparent phenotypic abnormalities. We now report that deletion of *Zfhx3* abrogates hypercapnia-induced suppression of antiviral and innate immune genes expression in murine MØs, and that myeloid *Zfhx3*-deficiency attenuates CO_2_-induced increases in viral replication, lung injury and mortality in mice infected with IAV. Our results define Zfhx3 as the first known component of a CO_2_ response pathway impacting the immune system and host defense in a mammalian system.

## Results

### Zfhx3 is expressed in mouse and human macrophages

Having previously stablished Zfh2 as a bona fide mediator of hypercapnic immune suppression in *Drosophila* ^19^, here we determined whether its ortholog, Zfhx3 (Atbf1), is expressed in mammalian MØs. Using specific antibodies, we demonstrated the presence of Zfhx3 protein by immunofluorescence microscopy (Figure 1A-D) and immunoblotting (Figure 1E) and confirmed the presence of Zfhx3 protein in both mouse (Figure 1A-C, E) and human MØs (Figure 1 D-E). Human MØs and BMDM both demonstrated a major band at ∼400 kD, corresponding to Zfhx3-A, and a minor band at ∼300 kD, corresponding to Zfhx3-B (Figure 1E). In addition, expression of *Zfhx3* mRNA in mouse AMØs was confirmed by qPCR (Supplemental Figure 1E).

**Figure 1:**
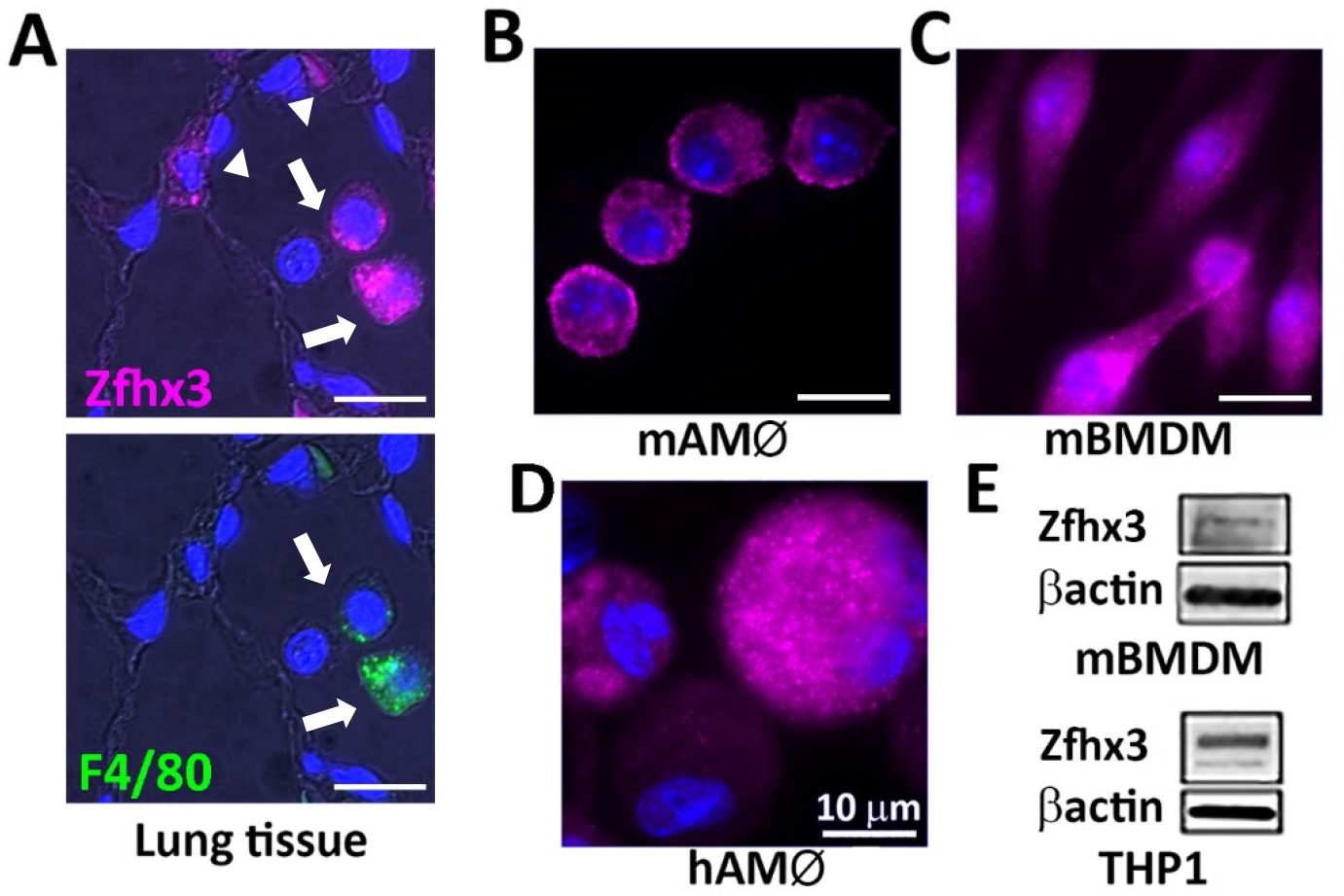
Mouse and human alveolar macrophages express Zfhx3. Mouse lung tissue (A), mouse AMØs (B), cultured mouse BMDM (C), and human AMØs (D) were fixed and stained with specific anti-Zfhx3 antibody (magenta). Lung tissue in (A) was double stained with anti-F4/80 (green) to identify MØs (white arrows). In A (top panel), alveolar epithelial cells (arrow heads) also stain for Zfhx3. Nuclei stained with DAPI (blue). Scale bars: 10 µm. Immunoblot of mouse BMDM and human MØs (THP1 cells) for Zfhx3 and βactin (E).

### Zfhx3 expression is markedly reduced in alveolar macrophages from *Zfhx3*^*fl/fl*^*LyzM*^*Cre*^ mice

To determine the role of Zfhx3 in the response to hypercapnia, we sought to study a mouse that was genetically deficient in *Zfhx3*. However, global homozygous *Zfhx3* deficiency is a lethal mutation and heterozygosity results in pre-weaning mortality ^24^. For this reason, and because our previous findings strongly implicate the MØ as a key cellular target of hypercapnic immunosuppression ^18^, we crossed a *Zfhx3*^*fl/fl*^ mice ^24^ with *LysM*^*Cre*^ mice (*Lyz2*^*tm1(cre)Ifo*^, Jackson Labs) to generate mice with homozygous *Zfhx3* deficiency in the myeloid lineage. The myeloid-selective *Zfhx3*^*-/-*^ mutant (for simplicity, referred to hereafter as *Zfhx3*^*-/-*^) is fully viable, grows normally, and exhibits no apparent phenotypic abnormalities in the unchallenged state. Immunofluorescence microscopy shows that while AMØs obtained by bronchoalveolar lavage (BAL) and lung tissue from *Zfhx3*^*fl/fl*^ controls (referred to hereafter as *Zfhx3*^*+/+*^), stain strongly for Zfhx3, the protein is undetectable in AMØs from Zfhx3^-/-^ mice (Supplemental Figure 1A and 1B). Likewise, Zfhx3 protein is highly expressed in BMDM from *Zfhx3*^*+/+*^ mice, but undetectable by immunofluorescence or immunoblot in BMDM from *Zfhx3*^*-/-*^ mice (Supplemental Figure 1C and 1D). Also, *Zfhx3* mRNA is >95% decreased by qPCR in BMDM from *Zfhx3*^*-/-*^ mice compared to Zfhx3^*+/+*^ controls (Supplemental Figure 1E). Besides AMØs, Zfhx3 is expressed in the lung in surfactant protein C (SPC)-positive AT2 epithelial cells (Supplemental Figure 1B). Because a small percentage of lung epithelial cells express LysM and can be targeted by *LysM-Cre* constructs ^29^, we quantified Zfhx3 protein in SPC-positive AT2 cells in lung tissue from *Zfhx3*^*+/+*^ and *Zfhx3*^*-/-*^ mice. This analysis showed a small but non-significant decrease of Zfhx3 expression, measured as corrected total cell fluorescence (CTCF), in AT2 cells from *Zfhx3*^*-/-*^ as compared to *Zfhx3*^*+/+*^ mice (Supplemental Figure 1F), indicating that *LysM*^*Cre*^-mediated *Zfhx3* knockdown in the lung was indeed highly selective for the myeloid lineage, as intended.

### Hypercapnia selectively alters gene expression in alveolar macrophages, resulting in down-regulation of innate immune and antiviral pathways

To assess the impact of elevated CO_2_ on global gene expression in AMØs, mice were exposed to ambient air or normoxic hypercapnia (HC, 10% CO_2_/21% O_2_) for 7 days, which we previously showed increases arterial PCO_2_ to ∼75 mm Hg, as compared to ∼40 mm Hg in air-breathing animals ^17^. After exposure to 10% CO_2_ or air as control for 7 days, lungs were harvested, and flow-sorted AMØs were subjected to bulk RNA-sequencing. Global gene expression analysis revealed that hypercapnia significantly modified the expression of 878 genes ≥ ±1.4 fold (expressed as log_2_ [fold change]) with an adjusted P value ≤ 0.05, including 619 downregulated and 259 upregulated genes (Figure 2A). The volcano plot (Figure 2B) shows that the fold change for a large proportion of the downregulated genes was much greater than that for the upregulated genes. Differential gene expression was consistent in AMØs from replicate mice, as represented in the heat map with K-means clustering (Figure 2C).

**Figure 2:**
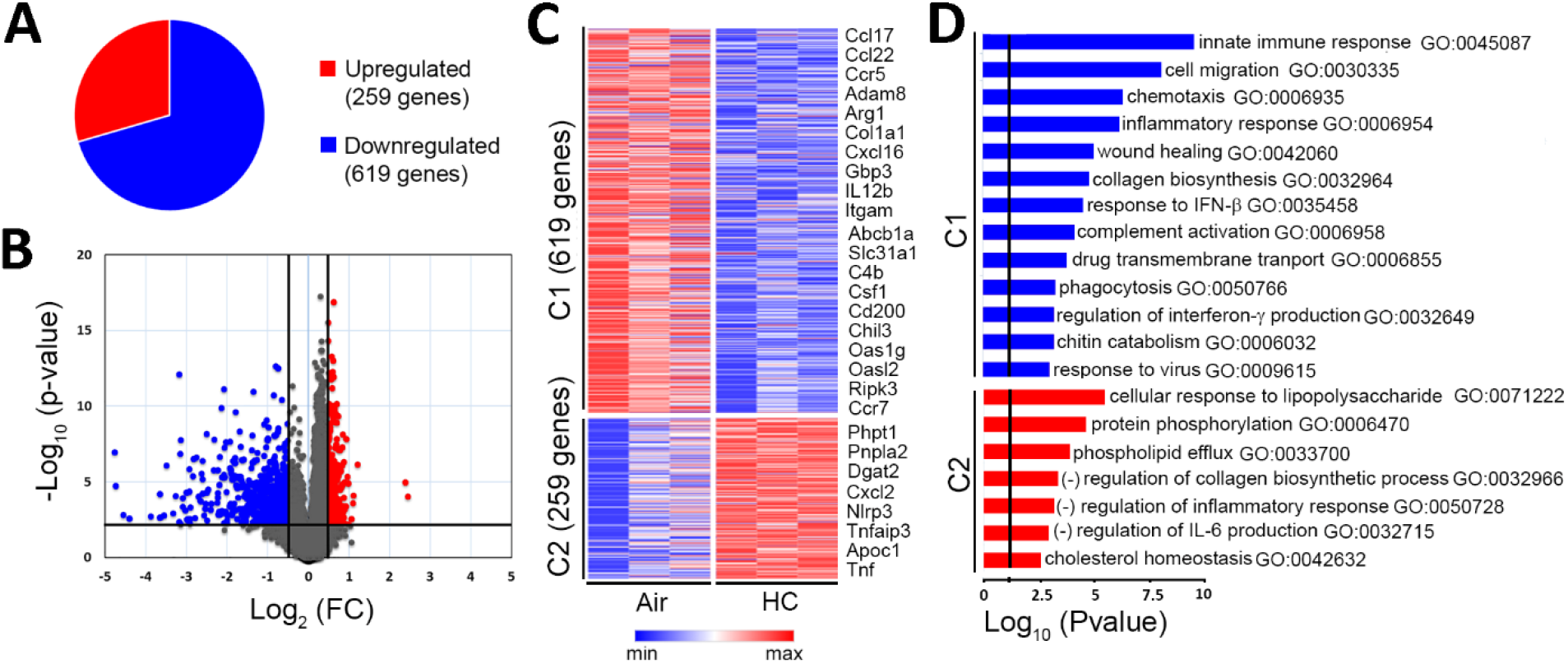
Hypercapnia alters gene transcription, including down-regulation of innate immune and antiviral genes, in alveolar macrophages. RNA-seq analysis was performed in AMØs obtained from mice exposed to room air or 10% CO_2_ (hypercapnia, HC) for 7 days and isolated by flow cytometry. Pie chart indicating proportion of genes downregulated or upregulated by hypercapnia (A). Volcano plot showing statistical significance (−log_10_ [P value]) plotted against log_2_ fold change for hypercapnia vs air (B). Plot indicates significantly upregulated genes (log_2_ [fold change] ≥+0.5, adjusted P value ≤0.05) in red and downregulated genes (log_2_ [fold change] ≤−0.5, adjusted P value ≤0.05) in blue. K-means clustering was performed on differentially expressed genes and is presented as a heatmap of cluster 1 (C1) and cluster 2 (C2) (C). Bars represent the top GO biological processes of downregulated (blue, C1) and upregulated (red, C2) genes induced by elevated CO_2_ (D).

Gene Ontology (GO) analysis shows that hypercapnia downregulated multiple processes related to innate immunity and response to viral infection in AMØs (Figure 2D and Supplemental Figure 2A). Genes whose change in expression maps to antiviral processes downregulated for hypercapnia include Gbp6, Iigp1, Pyhin1, Bcl3, Clec5a and Stmn1, and the interferon (IFN)-stimulated antiviral effector Oasl2. Additional innate immune response genes downregulated by elevated CO_2_ relate to inflammatory responses (Adam8, Ccl22, Ccr5, Ccr7, Irg1, Ptgs2), chemotaxis (C3ar1, Ccl17), complement activation (C1q, Cfh, Hc, Serping1) and phagocytosis (Mfge8, Sftpa1, Slc11a1). Other hypercapnia-regulated genes map to processes downregulated by elevated CO_2_ that could also influence outcomes of infection; these include cell migration (Csf1, Cyr61, Hbegf, Mmp14, Pdgfr), wound healing (Dcbld2, Hpse, Mmp12, Mustn1, Pdgfb, Pdgfra, Slc11a1, Sparc, Timp1), collagen biosynthesis (Arg1, Col1a1, Col3a1, Serpinh1) and chitin catabolism (Chil3 and Chil4). Notable among the GO biological processes upregulated by hypercapnia are negative response to inflammation (Nlrp3, Nt5e, Tnfaip3, Zfp36), and to IL-6 (Tnf, Tnfaip3, Zc3h12a) (Figure 2D and Supplemental Figure 2B). Supplemental Figure 2A and 2B shows that many CO_2_-regulated genes map to gene networks shared by more than one GO process downregulated or upregulated by hypercapnia. In summary, the changes in AMØ gene expression resulting from exposure to elevated CO_2_ suggest mechanisms by which hypercapnia would be expected to increase susceptibility to and/or worsen the outcome of infections due to viruses and other pathogens.

### *Zfhx3* deficiency abrogates hypercapnia-induced changes in expression of innate immune and inflammatory pathway genes in alveolar macrophages

Previously, we showed that knockdown of *zfh2* blocked hypercapnic suppression of multiple antimicrobial peptides in *Drosophila* ^*19*^. Here, we were interested in determining the impact of myeloid *Zfhx3* deficiency on hypercapnia-induced alterations in expression of immune, inflammatory, and other genes. Thus, we exposed *Zfhx3*^*-/-*^ and *Zfhx3*^*+/+*^ mice to normoxic hypercapnia (10% CO_2_/21% O_2_) or ambient air for 7 days, flow sorted AMØs, performed RNA sequencing, as in previous experiment. The heat map in Figure 3A shows that *Zfhx3* deficiency abrogated hypercapnic downregulation of 124 genes (cluster 1, C1) and CO_*2*_-induced upregulation of 91 genes (cluster 2, C2), representing subsets of 619 genes downregulated and 259 genes upregulated by hypercapnia in *Zfhx3*^*+/+*^ AMØs. GO analysis shows that the biologic processes upregulated under hypercapnic conditions in *Zfhx3*^*-/-*^ as compared to *Zfhx3*^*+/+*^ AMØs include innate immunity and inflammatory responses, complement activation and chemotaxis (Figure 3B). Genes whose change in expression maps to these processes include Anpep, C1q, C3ar1, C4b, Ccl17, Cd200, Cd36, Cfh, Cxcl16, Ear11, Igf1, Itgam, Lgmn, Lrrk2, Msr1, Raet1d, Ripk3, Thbs1, Treml2, C4b, Chil3, Chil4, Cxcl15, Pla2g7, Slc11a1, Thbs1. Biologic processes downregulated by hypercapnia in *Zfhx3*^*-/-*^ as compared to *Zfhx3*^*+/+*^ AMØs include response to LPS and cellular response to cAMP, including the genes Ccrn4l, Il1rn, Nfkbia, Tnf, Egr3 and Gpd1. Multiple genes that were differentially expressed in *Zfhx3*^*-/-*^ as compared to *Zfhx3*^*+/+*^ AMØs in under hypercapnic conditions map to more than one process (Supplemental Figure 3A and 3B). The foregoing results indicate that myeloid *Zfhx3* deficiency protects AMØs from CO_2_-induced changes in expression of specific sets of genes in a manner that might be expected to prevent or attenuate hypercapnic immunosuppression.

**Figure 3:**
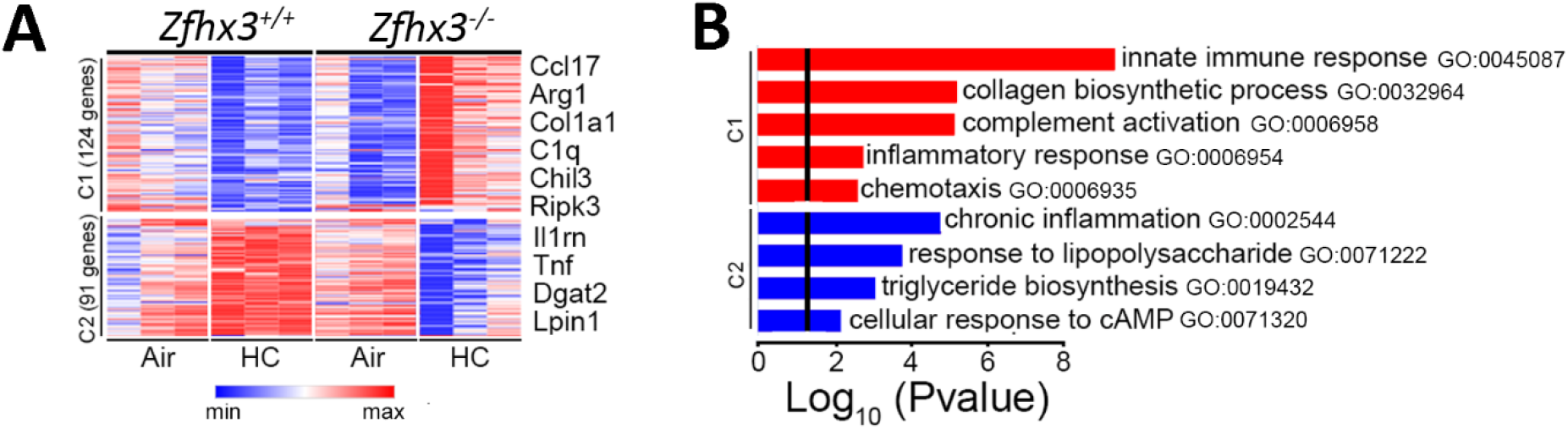
Myeloid *Zfhx3* deficiency abrogates hypercapnia-induced changes in expression of innate immune and inflammatory pathway genes. *Zfhx3*^*+/+*^ and *Zfhx3*^*-/-*^ mice were exposed to room air or 10% CO_2_ (hypercapnia, HC) for 7 days. AMØs from those mice were isolated by flow cytometry assessed by RNAseq analysis. K-means clustering performed on differentially expressed genes and presented as a heatmap (A). Bars represent the top GO biological processes of each cluster upregulated by HC and reversed in *Zfhx3*^-/-^ AMØs (blue) or downregulated by high CO_2_ and reversed in *Zfhx3*^-/-^ AMØs (red) (B).

### Myeloid *Zfhx3* deficiency protects against hypercapnia-induced increases in lung injury and mortality in mice infected with IAV

To determine the importance of Zfhx3 in CO_2_-induced suppression of antiviral host defense, we infected *Zfhx3*^*-/-*^ and *Zfhx3*^*+/+*^ mice with IAV in the absence or the presence of hypercapnia. Prior to infection, to avoid the confound effects of transient acidosis resulting from chronic hypercapnia exposure, mice were exposed to ambient air or normoxic hypercapnia (10% CO_2_/21% O_2_) for 3 days, which we previously showed allows for maximal renal compensation of respiratory acidosis, resulting in an arterial pH of ∼7.3, close to that of air-breathing controls ^17^. Mice were then inoculated with IAV (A/WSN/1933) at 3 or 30 pfu and maintained in air or hypercapnia, respectively until sacrifice at 4 or 7 dpi, or monitored until death or recovery. As previously reported in wild-type mice ^18^, exposure to elevated CO_2_ worsened IAV-induced histologic lung injury in *Zfhx3*^*+/+*^ mice, both following infection with 30 (Figure 4A and 4B) and 3 (Supplemental Figure 4A) pfu per animal. By contrast, *Zfhx3*^*-/-*^ mice were protected against the increase in IAV-induced lung injury caused by hypercapnia as shown in representative images (Figure 4A and Supplemental Figure 4A) and by histopathologic score (Figure 4B). We assessed the effect of myeloid *Zfhx3* deficiency on the mortality of IAV infection in mice inoculated with 3 pfu per animal, at which dose all air breathing *Zfhx3*^*+/+*^ mice survived, while those exposed to hypercapnia lost more weight (Figure 4C) and exhibited 100% mortality by 11 dpi (Figure 4D). On the other hand, *Zfhx3*^*-/-*^ mice exposed to hypercapnia and infected with IAV survived longer than *Zfhx3*^*+/+*^ mice, and mortality of IAV infection in the *Zfhx3*^*-/-*^ mice was reduced to 73% (Figure 4D). Interestingly, 10% CO_2_-exposed *Zfhx3*^*+/+*^ and *Zfhx3*^*-/-*^ mice lost similar amount of weight after IAV infection (Figure 4C), suggesting that the protective effect myeloid Zfhx3 deficiency on the outcome of IAV infection in the setting of hypercapnia was not due to improved intake of food and water in *Zfhx3*^*-/-*^ mice. Taken together, these results establish *Zfhx3* as a bona fide mediator of CO_2_-induced immunosuppression in a mammalian system.

**Figure 4:**
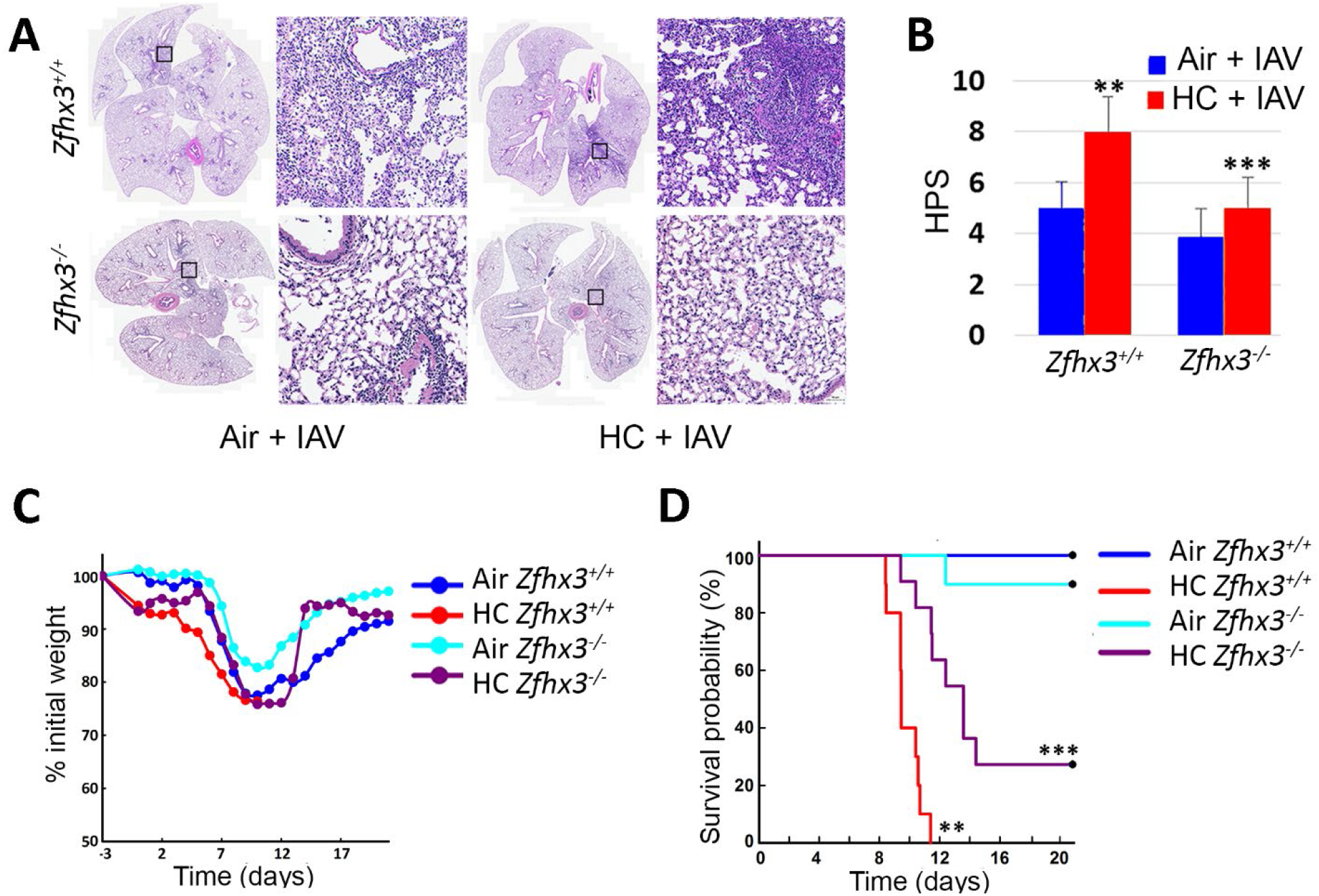
Myeloid *Zfhx3* deficiency protects against hypercapnia-induced increases in lung injury and mortality in IAV-infected mice. *Zfhx3*^*+/+*^ and *Zfhx3*^*-/-*^ mice pre-exposed to normoxic hypercapnia (10% CO_2_/21% O_2_, HC) for 3 days, or air as control, then infected intratracheally with 30 (A-B), or 3 (C-D) pfu IAV (A/WSN/33) per animal; N = 6-10 per group. Lungs from IAV-infected mice harvested 4 dpi sectioned and stained with H&E (A) and images were assessed blindly to determine histophatologic scores (HPS) for lung injury (B). Body weight changes over time (C) and Kaplan-Meier plots show survival and analyzed by log-rank test (D) after infection with 3 pfu IAV. **P<0.05 vs Air + IAV *Zfhx3*^*+/+*^, ***P<0.05 vs HC + IAV *Zfhx3*^*+/+*^.

### Myeloid *Zfhx3* deficiency prevents hypercapnia-induced increases in viral replication and suppression of antiviral gene and protein expression in IAV-infected mice

To understand mechanisms by which myeloid *Zfhx3* deficiency protects against the adverse impact of hypercapnia on IAV-induced lung injury and mortality, we assessed viral loads and expression of viral and antiviral genes and proteins in lung tissue and AMØs from IAV-infected *Zfhx3*^*+/+*^ and *Zfhx3*^*-/-*^ mice. First, we found that while exposure to 10% CO_2_ increased the amount of live virus recoverable from the lungs of *Zfhx3*^*+/+*^ mice 4 dpi by more than 2-fold, the hypercapnia-induced increase in viral load was largely blocked in *Zfhx3*^*-/-*^ mice (Figure 5A). Likewise, 10% CO_2_ exposure increased expression of the viral proteins NS1 and M2 in the lungs of IAV-infected *Zfhx3*^*+/+*^ mice, but this effect of hypercapnia was blocked in *Zfhx3*^*-/-*^ mice (Figure 5B and Supplemental Figure 4B and 4D). Similarly, hypercapnia increased NS1 mRNA expression in lung tissue of *Zfhx3*^*+/+*^ mice, as assessed by RNAscope®, and this was largely attenuated in the lungs of CO_2_-exposed *Zfhx3*^*-/-*^ mice (Figure 5C, left panel). Conversely, exposure to hypercapnia blunted or suppressed expression of mRNA and protein for IFN-β and the IFN-stimulated antiviral effectors, viperin and Oas1, in IAV-infected *Zfhx3*^*+/+*^ mice, whereas mRNA and protein expression of these antiviral factors was increased in the lungs of CO_2_-exposed *Zfhx3*^*-/-*^ mice (Figure 5C, middle and right panels, and Supplemental Figure 4C and 4D).

**Figure 5:**
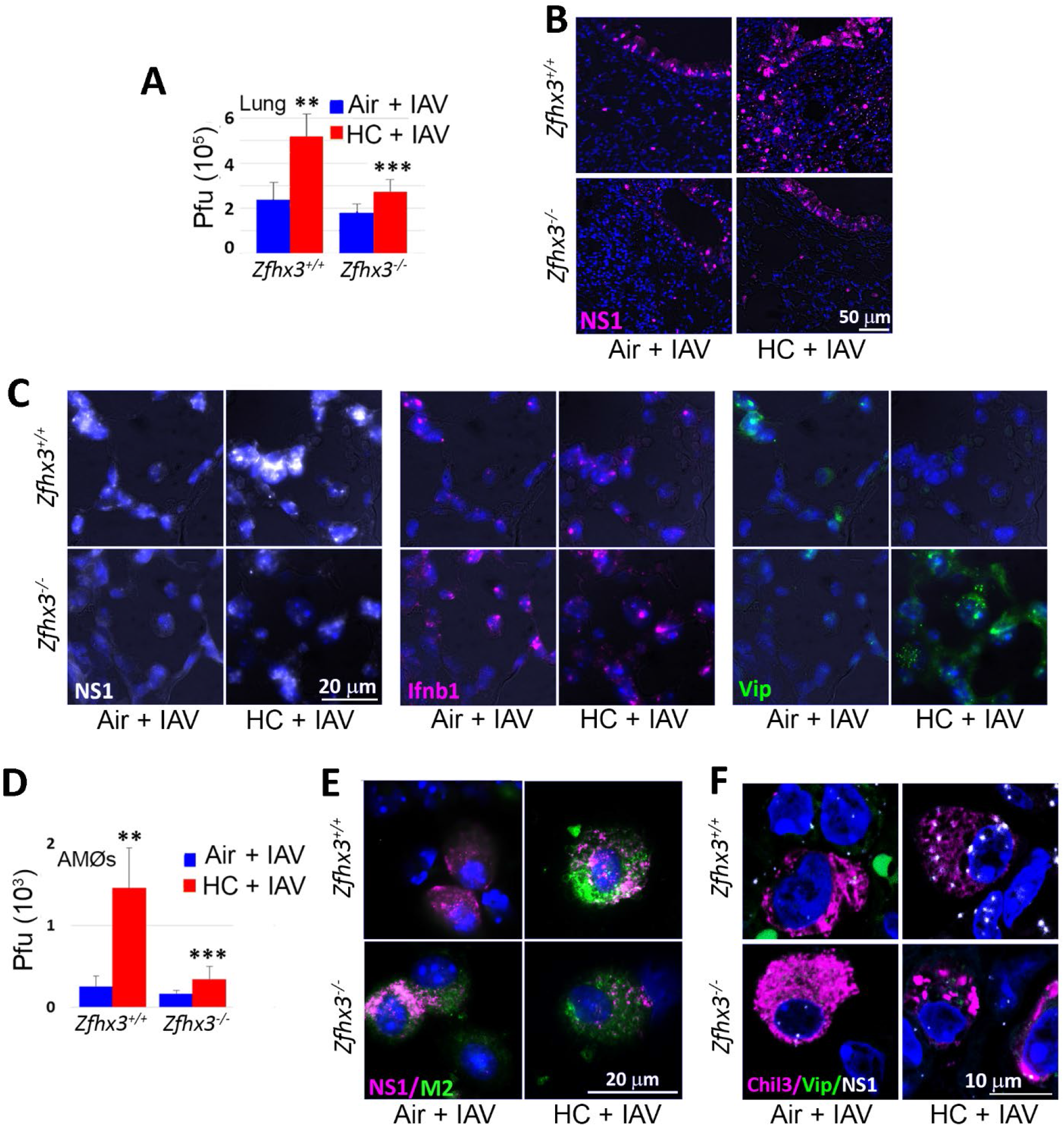
Myeloid *Zfhx3* deficiency increases viral replication and prevents hypercapnia-induced suppression of antiviral gene and protein expression in IAV-infected mice. *Zfhx3*^*+/+*^ and *Zfhx3*^*-/-*^ mice pre-exposed to normoxic hypercapnia (10% CO_2_/21% O_2_, HC) for 3 days, or air as control, then infected intratracheally with 30 (A-B, D-F) or 300 (C) pfu IAV (A/WSN/33) per animal; N = 6-10 per group. Viral titers in homogenized lung tissue determined by plaque assay at 4 dpi (A). Expression of viral NS1 (magenta) assessed in lung tissue sections from mice sacrificed 4 dpi (B). Ifnb1, viperin and viral NS1 transcript expression in lung tissue sections from mice infected with IAV 300 pfu 1 dpi was detected by RNAscope (C). AMØs from IAV-infected mice were obtained by BAL 1 dpi and cultured under normocapnic (5% CO_2_/95% air, NC) or hypercapnic (15% CO_2_/21% O_2_/64% N_2_, HC) conditions for 18 h, after which viral titers in culture supernatants were determined by plaque assay (D) or assessed for viral NS1 (magenta) and M2 (green) protein expression by immunofluorescence (E). Viperin and NS1 were also assessed by RNAscope, in combination with Chil3 (AMØ marker) by immunofluorescence (F). Nuclei were stained with DAPI (blue). **P<0.05 vs Air + IAV *Zfhx3*^*+/+*^, ***P<0.05 vs HC + IAV *Zfhx3*^*+/+*^.

To determine the effects of myeloid *Zfhx3* deficiency on the response of lung MØs specifically, we studied AMØs isolated from *Zfhx3*^*+/+*^ and *Zfhx3*^*-/-*^ mice exposed to 10% CO_2_ or ambient air and infected with IAV (300 pfu), as described above. AMØs were obtained by BAL 1 dpi, washed, and cultured under normocapnic (5% CO_2_/95% air, NC) or hypercapnic (15% CO_2_/21% O_2_/64% N_2_, HC) conditions for 18 h prior to analysis. Figure 5D shows that exposure of *Zfhx3*^*+/+*^ AMØs to hypercapnia as compared to air/NC resulted in a 6-fold increase in recovery of viable virus, strongly suggesting active viral replication in these cells in the setting of elevated CO_2_. As in the homogenized lung, the hypercapnia-induced increase in viral load was almost fully blocked in AMØs from *Zfhx3*^*-/-*^ mice (Figure 5D). Also, mirroring the finding in lung tissue, hypercapnia increased expression of viral NS1 and M2 protein (Figure 5E) and NS1 mRNA (Figure 5F) and reduced viperin mRNA expression (Figure 5F) in AMØs from *Zfhx3*^*+/+*^ mice. These effects of elevated CO_2_ were blocked in AMØs from *Zfhx3*^*-/-*^ mice (Figure 5E and 5F). Myeloid *Zfhx3* deficiency also abrogated the increase of viral proteins NP, NS1 and M2 (Supplemental Figure 5A-5D) and suppression of antiviral protein expression of Oas1 and viperin (Supplemental Figure 5C-D) caused by hypercapnia in IAV-infected mouse BMDM.

Exposure of mice to elevated CO_2_ increased viral protein expression not only in AMØs, but also in lung parenchymal cells, including AT2 and bronchial epithelial cells ^18^, Figure 5B and Supplemental Figure 4B and 4D). Of note, myeloid *Zfhx3* deficiency not only blocked the hypercapnia-induced increase in expression of viral gene products and suppression of host antiviral factors in MØ, but also blunted the increase in viral NS1 and M2 and the decrease in the antiviral effector viperin in airway and alveolar cells of CO_2_-exposed mice (Figure 5B and Supplemental Figure 4B and 4D). This suggest myeloid *Zfhx3* deficiency protects against hypercapnic suppression of antiviral host defense in vivo not only by enhancing the resistance of MØs to viral infection, but also by reducing viral infection and/or replication in lung epithelial cells.

## Discussion

In the present work, we show that Zfhx3, mouse ortholog of zfh2, identified previously as a CO_2_ mediator of CO_2_ effects in *Drosophila*, is highly expressed in mouse and human MØs. We document that exposure of mice to 10% CO_2_ selectively alters gene expression in AMØs, resulting in prominent downregulation of innate immune and antiviral gene programs, and that knockout of *Zfhx3* in the myeloid lineage prevents hypercapnic downregulation of innate immune pathways and also blocks CO_2_-induced changes in other gene programs. Moreover, we found that myeloid *Zfhx3*^*-/-*^ mice were protected from hypercapnia-induced increases in viral replication, inflammatory lung injury and mortality following IAV infection. Myeloid *Zfhx3* deficiency blocked hypercapnic suppression of IFN-β and antiviral effector gene and protein expression and the CO_2_-induced increase in viral gene and protein expression caused by hypercapnia in IAV-infected *Zfhx3*^*+/+*^ mice. These findings establish *Zfhx3* as the first known mediator of CO_2_-induced immunosuppression in a mammalian system.

We chose to investigate the role of Zfhx3 as a potential mediator of CO_2_ effects on the immune system in mice because of its homology to *Drosophila zfh2*, which we previously identified as a mediator of CO_2_-induced suppression of host defense against bacterial infection in the fly ^19^. While full length human Zfhx3 (Zfhx3-A) has 23 zinc fingers and 4 homeodomains CO_2_-induced, as compared to 16 zinc fingers and 3 homeodomains in zfh2 ^30^, key regions of the two proteins are highly homologous: homeodomains I, II and III of Zfhx3 are 77, 69 and 61% identical to the corresponding homeodomains in zfh2; 13 of the zinc fingers in Zfhx3 are from 22 to 89% identical to those in zfh2; and all of these domains in the two proteins are co-linearly arranged ^20^. Knockdown of *zfh2* abrogated CO_2_-induced suppression of multiple antimicrobial peptides in cultured *Drosophila* S2* phagocytes ^19^, like the effect of *Zfhx3* deletion on hypercapnic suppression of innate immune and antiviral genes in mouse and human MØs. Further, in vivo knockdown of *zfh2* in the immune responsive fat body protected *Drosophila* from the increase in mortality of bacterial infection caused by elevated CO_2 19_, similar to the effect of myeloid *Zfhx3* deficiency on the hypercapnia-induced increase in mortality of IAV infection in mice. The high degree of homology between zfh2 and Zfhx3, and the parallels between the impact of knocking down or knocking out the respective genes in *Drosophila* and mice strongly suggest that the role of Zfhx3 as a mediator of CO_2_ effects on the immune system has been evolutionarily conserved.

Its large size and multiplicity of homeodomains and zinc fingers suggests the possibility of numerous protein-DNA and protein-protein interactions through which Zfhx3 may mediate CO_2_ effects on gene and protein expression. Several different Zfhx3-DNA and -protein interactions have been elucidated in previous studies unrelated CO_2_ or the immune system. ATBF1, the alternate name for Zfhx3, derives from the observation that the protein negatively regulates transcription of alpha-fetoprotein (AFP) by binding to AT-rich elements in the AFP promoter and enhancer ^31^. Alternately, ATBF1 was shown to bind to a leucine zipper motif of the oncoprotein v-Myb and thereby inhibit its oncogenic transcriptional activity ^32^. Also, ATBF1 was found to bind to and inhibit transcriptional activity of the estrogen receptor by blocking its ability to interact with the steroid receptor co-activator AIB1 in breast cancer ^33^. Another study showed that upon stimulation with TGF-β, ATBF1 bound the transcription factor RUNX3 through an undefined interaction, resulting in co-translocation to the nucleus and synergistic upregulation of *p21*^*Waf1/Cip1*^ promoter activity in gastric cancer cells ^34^. These examples highlight the possibility that Zfhx3 may mediate effects of hypercapnia on transcription of immune and antiviral gene by direct binding to their promoters and/or other regulatory DNA elements, or by interacting with other transcription factors or upstream components of a putative CO_2_-triggered signaling pathway.

The fact that deletion of *Zfhx3* in the myeloid lineage attenuates the hypercapnia-induced increase in lung injury and mortality of IAV infection underscores the importance of lung MØs as a key target of elevated CO_2_ in suppressing antiviral host defense. The critical role of AMØs in host defense against influenza viruses has been established by studies in several species showing that ablation of AMØs with intratracheal clodronate or by genetic strategies increases the mortality of IAV infection ^25-28^. The protective effects of AMØs involve both robust type I IFN antiviral signaling and modulation of the inflammatory response to viral infection. While IAV can infect mouse and human AMØs, most studies show that infection is largely abortive, resulting in release of minimal amounts of infectious new virions ^35^. However, elevated CO_2_ suppresses the AMØ antiviral response, such that BAL AMØs obtained one day after intratracheal inoculation of hypercapnic mice with IAV expressed increased amounts of viral NS1 and M2 and released 6-fold more viable virus than AMØs from mice breathing ambient air (Figure 5 D-F). The increase in mRNA and protein expression of NS1, a nonstructural viral gene product that inhibits transcription of antiviral host genes and blocks the activity of IFN-stimulated antiviral effectors ^36^, is indicative of active viral replication in AMØs from hypercapnic mice. Importantly, myeloid *Zfhx3* deficiency prevented hypercapnia from suppressing antiviral gene and protein expression, and from increasing viral NS1 and M2 expression and viral replication in AMØs from IAV-infected mice. Of note, *Zfhx3* deficiency also reduced viral protein expression in lung epithelial cells (Figure 5B and 5C and Supplemental Figure 4B and 4D) and attenuated the hypercapnia-induced increases in lung injury and mortality due to IAV infection in *Zfhx3*^*+/+*^ mice. Because LysM is expressed in a fraction of AT2 cells, *LysM*^*Cre*^-mediated recombination may have deleted *Zfhx3* in this AT2 cell subset, and it is possible that this could contribute to protection from the adverse effects of hypercapnia in *Zfhx3*^*-/-*^ mice. However, Zfhx3 protein expression was not decreased in AT2 cells of *Zfhx3*^*-/-*^ mice as compared to *Zfhx3*^*+/+*^ mice, in contrast to the >95% reduction in Zfhx3 mRNA and protein in *Zfhx3*^*-/-*^ MØs. This suggests that preventing CO_2_-induced suppression of the antiviral response through *Zfhx3* deficiency in MØs (and other myeloid cells) also decreases viral replication in lung epithelial cells and that selective myeloid Zfhx3 deficiency is sufficient to reduce IAV-induced lung injury and mortality in the setting of hypercapnia.

While our findings emphasize the importance of Zfhx3 as a mediator of CO_2_ effects on AMØs, we also demonstrate expression of Zfhx3 mRNA and protein in alveolar epithelial cells (Supplemental Figure 1B and 1D). Moreover, single-cell RNA sequencing documents *Zfhx3* expression in a broad array of cell types during human lung development ^37^, and specifically in AMØs, alveolar epithelial cells, ciliated and mucus-secreting airway epithelial cells, fibroblasts, smooth muscle cells, adventitial cells and others in adult mouse and human lungs ^38,39^. Widespread expression among many, but not all, cell types during development and in the adult lung suggests that Zfhx3 has multiple functions, as would be expected for such a large and complex protein. Further, Zfhx3 may mediate CO_2_-induced changes in gene expression in lung cells other than AMØs that also impact antiviral host defense. Indeed, we previously showed that hypercapnia selectively alters global gene expression in human bronchial epithelial cells, including downregulation of innate immune pathways ^40^. This, plus the observation that myeloid *Zfhx3* deficiency only partially protects mice from the increase in mortality of IAV infection caused by hypercapnia, suggests that future studies should examine the role of Zfhx3 in responses to elevated CO_2_ in other cells, particularly airway and alveolar epithelial cells, in the context of viral infection.

The molecular mechanism by which elevated CO_2_ impacts Zfhx3 remains to be elucidated. This is unlikely to be an effect of altered pH fort two reasons. First, in previous studies with cultured cells, we have documented that hypercapnia-induced changes in gene expression are not a function of extracellular or intracellular acidosis ^14,16,18^. Second, in the current investigation, as in our previous in vivo studies ^17,18^, mice were exposed to 10% CO_2_ for 3 days, allowing for maximal renal compensation of respiratory acidosis and return of pH to near normal, prior to IAV or bacterial infection. One possible mechanism by which molecular CO_2_ might trigger alterations in gene expression is by carbamylation of regulatory proteins. Carbamylation by reaction of CO_2_ with the amino group of lysine residues represents a recently described post-translational modification that can alter protein function ^41,42^. Whether Zfhx3 or upstream components of a putative CO_2_ signaling pathway undergo differential carbamylation at normal vs elevated levels of PCO_2_ is another direction for future investigation.

In summary, in the present work we show that knockout of Zfhx3 in the myeloid lineage blocks hypercapnia-induced downregulation of antiviral and innate immune gene expression and protects mice from CO_2_-induced increases in viral replication, inflammatory lung injury and mortality when infected with IAV. These results establish Zfhx3 as the first known mediator of CO_2_ effects on the immune system mammals and suggest that this function has been conserved across evolution from *Drosophila* zfh2. Further studies are needed to define additional components of the pathway(s) through which elevated CO_2_ signals, including the CO_2_ sensor and the mechanism by which CO_2_ is sensed. This avenue of investigation has the potential to identify new targets for pharmacologic intervention to mitigate hypercapnia-induced immunosuppression, with the goal of improving outcomes of infection in patients with severe acute and chronic lung diseases.

## Materials and Methods

### Mice

*Zfhx3*^*fl/fl*^*LyzM*^*Cre*^ mice lacking *Zfhx3* in the myeloid lineage (*Zfhx3*^*-/-*^ mice) was generated by crossing C57Bl/6 *Zfhx3*^*fl/fl*^ mice ^24^ with *LyzM*^*Cre*^ mice (B6.129P2-Lyz2^tm1(cre)Ifo^/J, The Jackson Laboratory). Myeloid *Zfhx3*^-/-^ mice survive, grow, and appear phenotypically normal up to > 1 year of age. In all experiments, *Zfhx3*^*fl/fl*^ mice (*Zfhx3*^*+/+*^ mice) were used as controls. Given that we observed no sex-related differences in responses to hypercapnia or outcomes of IAV infection in previous work with wild type mice and in preliminary experiments with *Zfhx3*^*-/-*^ mice, mixed groups of male and female animals were used in all experiments. Mice aged 6 to 12-week were used. The work was performed according to a protocol approved by the Institutional Animal Care and Use Committee of Northwestern University, and compliant with to National Institutes of Health guidelines for the use of rodents.

### Murine CO_2_ exposure

*Zfhx3*^*+/+*^ and *Zfhx3*^*-/-*^ mice were exposed to normoxic hypercapnia (10% CO_2_/ 21% O_2_/ 69% N_2_) in a BioSpherix A environmental chamber (BioSpherix). O_2_ and CO_2_ concentrations in the chamber were maintained at the indicated levels using ProOx C21 O_2_ and CO_2_ controllers (BioSpherix). Age-and genotype-matched mice, simultaneously maintained in air, served as controls in all experiments.

### In vivo influenza virus infection

Mice pre-exposed to ambient air or hypercapnia for 3 days were anesthetized with isoflurane, intubated with a 20-gauge Angiocath™ catheter, and inoculated intratracheally with mouse-adapted IAV strain A/WSN/33, H1N1 gift of Robert Lamb, Ph.D., Sc.D., Northwestern University, Evanston, IL, or PBS as control, then maintained under air or hypercapnia exposure conditions, as previously described ^18^. Depending of the experiment, mice were infected with either 3, 30, or 300 pfu/animal in 50 µL of PBS.

### Clinical assessment of influenza A infection

After infection with IAV, mice were weighed daily and monitored every 8 h for signs of severe distress (slowed respiration, failure to respond to cage tapping, failure of grooming and/or fur ruffling). Mice that developed severe distress were considered moribund and sacrificed, and the deaths were recorded as IAV-induced mortality. Mice that died between monitoring episodes were also recorded as IAV-induced mortality.

### Cells

AMØs were isolated by BAL in mice anesthetized with ketamine and xylazine. Tracheotomy was performed and a 26-gauge catheter was inserted into the trachea and secured with vinyl suture. One milliliter of ice-cold PBS was instilled and withdrawn serially three times. BAL fluid was centrifuged, and cells were allowed to adhere to plastic, nonadherent cells were removed, resulting in an adherent AMØs population ≥98% pure. AMØs were cultured in RPMI 1640 with 10% FBS, 2 mM l-glutamine, 1 mM sodium pyruvate, 20 μM 2-ME, 100 U/ml penicillin, and 100 μg/ml streptomycin (RPMI media) ^14^. Cells were rested for 24 h to allow the transient proinflammatory profile of freshly isolated AMØs to subside prior experimentation ^43^. BAL cells were also centrifuged in a cytospin (1000 rpm for 5 min) and fixed in 4% PFA. For mice bone marrow derived MØs (BMDM), bone marrow was isolated from *Zfhx3*^*+/+*^or *Zfhx3*^*-/-*^ mice following a standard protocol ^44^. Cells were plated in 10 cm tissue culture plates at a density of 3 × 10^6^ cells/ plate. Cells were cultured with L929 cell supernatants/RPMI/FBS media to induce MØs differentiation. Media were changed every 3 days and BMDM were harvested by scraping on day 6, followed by plating in 24 well plates at a density of 8 × 10^5^ cells/well.

Human AMØs were obtained by BAL from the contralateral lung of subjects undergoing bronchoscopy for clinical diagnosis of noninfectious focal lung lesions ^14,45^ under a protocol approved by the Northwestern University Institutional Review Board and cultured as for mouse AMØs. Human monocytic leukemia THP-1 cells (ATCC) were cultured in RPMI media and differentiated to a MØ phenotype by exposure to 5 nM PMA for 48 h ^14^.

### Exposure of cells to normocapnia and hypercapnia

Normocapnia consisted of standard incubator atmosphere: humidified 5% CO_2_ (PCO_2_ 36 mmHg)/95% air, at 37°C. Hypercapnia consisted of 15% CO_2_ (PCO_2_ 108 mmHg)/21% O_2_/64% N_2_. Cells were exposed to hypercapnia in an environmental chamber (C-174, BioSpherix) contained within the same incubator where control cultures were simultaneously exposed to normocapnia. In all cases, cells were exposed to hypercapnia or maintained in normocapnia as control for 18 h prior to infection with IAV. All media were presaturated with 5% or 15% CO_2_ respectively before addition to the cells.

### In vitro influenza virus infection

Macrophages were infected with IAV A/WSN/1933(H1N1) at 2 MOI per cell, using a single-cycle protocol. Cells were incubated with IAV in a 5 % or 15% CO_2_ atmosphere at 37° C for 1h, during which time plates were rocked every 15 min to distribute the virus evenly and to keep the monolayer moist. Cells were then washed twice with PBS to remove excess of virus and fresh RPMI media was added to the plates.

### Immunofluorescence microscopy in tissue sections and cell cultures

Sections were deparaffinized with xylene, rehydrated by using graded ethanol, and subjected to heat-induced antigen retrieval with 10 mM of sodium citrate buffer (pH 6.0). After blocking with BSA 1% (wt/vol) in PBS, tissues were incubated overnight with anti-Zfhx3, anti-NS1 or anti-M2 antibodies. Then, sections were washed with PBS and Alexa–labeled secondary antibodies (1 μg/ml) were added. Co-labeling with F4/80 or Chil3 (MØ markers) and SPC (alveolar epithelial cell type II marker) was achieved by the addition of anti-F4/80, anti-Chil3 or anti-SPC antibodies respectively. After washing, Alexa-conjugated antibodies (1 μg/ml) were added, and sections were incubated for 1 h at room temperature.

Cells were fixed with 4% PFA, permeabilized with 0.1% Triton X100 for 5 min, blocked with BSA 1% (wt/vol) in PBS and incubated overnight with specific antibodies against host Zfhx3, OAS1, viperin and Chil3, and viral NS1, M2 and NP. F4/80 was used as macrophage marker. Then, sections were washed with PBS and Alexa-labeled secondary antibodies were added. Complete antibody information is provided in the Supplemental Table 1. Nuclei were visualized with DAPI and slides were mounted with Gel/Mount (Biomeda). Nonimmune mouse, rabbit or goat IgGs were used as a negative control and the staining was negative for all non-immune controls regardless of protocol. Fluorescent images were obtained using Axiovert 200M Fluorescence Microscope (Zeiss). Images were obtained using the same exposure time for all samples from a given experimental set. To avoid saturation, the exposure time was selected based on the most brightly stained sample and used for all other samples in the set. This approach results in equal subtraction of background autofluorescence from all images.

### Multiplexed in situ hybridization RNAscope® Assay combined with immunofluorescence on tissue sections

RNAscope Multiplex V2 Assay (ACD) was performed on paraffin-embedded 5-μm slices of lung tissue using mild digest times according to manufacturer instructions. Hybridization was detected with Opal dyes from Akoya Biosciences. The following RNAscope® Probes were used: Mm-*Rsad2* (viperin) with Opal dye 520, Mm-*Ifnb1*-C2 with Opal dye 570 and V-Influenza-H1N1-NS2NS1-C3 with Opal dye 620. The RNA-Protein Co-detection Ancillary kit was used to co-label RNA with viral M2 and mouse Chil3 proteins. Nucleus were stained with DAPI. Images were acquired using an Axiovert 200M Fluorescence Microscope (Zeiss) or a Nikon A1C Confocal Microscope at the Center for Advanced Microscopy at Northwestern University Feinberg School of Medicine. Final images were rendered using Fiji.

### Immunoblotting

The presence of indicated proteins in lung or cell homogenates were assessed by immunoblotting using the following Abs: anti-Zfhx3, anti-βactin and anti-OAS1. Complete antibody information is provided in Supplemental Table 1. Membranes were incubated with IRDye (1:10,000, LI-COR) Biosciences or HRP-conjugated (1:5000) secondary Abs for 1 h at room temperature. Signals were captured using the LI-COR Odyssey Fc Imager and analyzed with ImageStudio™ software (LI-COR).

### Quantitative real-time PCR

RNA was extracted using RNeasy Mini Kit (Qiagen) and reverse transcribed using an iScript cDNA Synthesis Kit (Bio-Rad). Amplification was performed using the CFX Connect Real-Time System (Bio-Rad) and TaqMan FAM-labeled primer/probes sets Mm01240016_m1 for *Zfhx3* and Mm00607939_s1 for *βactin. Zfhx3* gene expression was normalized to *βactin*. Relative expression was calculated by the comparative CT method (ΔΔCT) ^46^.

### Tissue preparation and flow cytometry

Tissue preparation for flow cytometry analysis and cell sorting was performed as previously described ^47^. Cells were stained with eFluor 506 (eBioscience) viability dyes, incubated with FcBlock (BD Bioscinces), and stained with fluorochrome-conjugated antibodies (antibodies, clones, fluorochromes, and manufacturers were described in detail in our previous publication ^47^. Data were acquired on a BD LSR II flow cytometer (for information regarding instrument configuration and antibody panels, see ^47^. Compensation, analysis, and visualization of the flow cytometry data were performed using FlowJo software (Tree Star). “Fluorescence minus one” controls were used when necessary to set up gates. Cell sorting was performed at Northwestern University RLHCCC Flow Cytometry core facility on SORP FACSAria III instrument (BD Bioscience) with the same optical configuration as for flow cytometry on the LSR II, using a 100-µm nozzle and a pressure of 40 psi.

### RNA Sequencing library preparation

RNA quality and quantity were measured using High Sensitivity RNA ScreenTape and the Agilent 4200 Tapestation System (Agilent Technologies). Briefly, mRNA was isolated from 50 ng of purified total RNA using oligo-dT beads (New England Biolabs, Inc). NEBNext Ultra(tm) RNA kit was used for full-length cDNA synthesis and library preparation. Libraries were pooled, denatured and diluted, resulting in a 2.0 pM DNA solution. PhiX control was spiked at 1%. Libraries were sequenced on an Illumina NextSeq 500 instrument (Illumina Inc) using NextSeq 500 High Output reagent kit (Illumina Inc) (1×75 cycles) with a target read depth of approximately 5-10 million aligned reads per sample.

### Gene expression profiling (RNA-seq) and bioinformatic analysis

Computation intensive analysis was performed using “Genomics Nodes” on Northwestern’s High Performance Computing Cluster, Quest (Northwestern IT and Research Computing). Reads were demultiplexed using bcl2fastc, quality was assessed using FastQC, reads were trimmed and aligned to mm10 reference genome using TopHat2, read counts were associated with genes using the GenomicRanges ^48^, and differential gene expression was assessed using edgeR ^49,50^ R/Bioconductor packages. Genes with less than one normalized read count across at least half of the samples were filtered from all analyses. Pearson correlation matrix and clustering heat maps were built using Morpheus software (https://software.broadinstitute.org/morpheus). Over representation analysis (ORA) of gene ontology (GO) terms from biological processes of all genes downregulated or upregulated by hypercapnia were separately analyzed using the Gene Ontology Analysis InnateDB tool ^51^ which utilizes a manually-curated knowledgebase of genes, proteins, interactions and signaling pathways involved in mammalian innate immune responses. Results from the Innate DB analysis were confirmed using GeneGo Metacore (Clariviate), a separately curated database and pathway analysis tool. RNA-seq data have been deposited to the National Center for Biotechnology Information (NCBI) Gene Expression Omnibus (GEO; http://www.ncbi.nlm.nih.gov/projects/geo) complied with MIAME standards (accession number GSE183922). From the GO biological term results, 5-6 processes of interest were selected. Interaction networks were constructed for each set of DEGs associated with these 5-6 terms using the GeneMANIA plug-in ^52^ for Cytoscape 3.8.0 software ^53^. GeneMANIA can find other genes that are related to the set of input genes and produce a functional association network based on their relationships, such as pathways, co-expression, co-localization, genetic interaction, physical interaction, shared protein domains and so on, based on the published literature.

### Lung histopathology

Mice were euthanized and lungs were perfused via the right ventricle with 10 ml HBSS with Ca^2+^ and Mg^2+^. A 20-gauge angiocatheter was inserted into the trachea, and secured with a suture, the heart and lungs were removed en bloc, and lungs were inflated with up to 0.8 ml of formalin at a pressure not exceeding 16 cm H_2_O and fixed overnight at 4°C. Tissues were embedded in paraffin, sectioned (4 μm thickness), deparaffinized, and stained with H&E by the Northwestern University Mouse Histology and Phenotyping Laboratory. Images of lungs were obtained using TissueFAXS PLUS Scanning System (TissueGnostics) at the Northwestern University Center for Advanced Microscopy. Serial images were stitched into a high-resolution macroscopic montage. The severity of inflammatory lung injury was evaluated by observers blinded to experimental groups using an histopathologic score (HPS) that assigns values of 0 to 26 (least to most severe) based on assessment of quantity and quality of peribronchial inflammatory infiltrates, luminal exudates, perivascular and parenchymal infiltrates and thickening of the membrane wall, as described previously ^54,55^. This scoring system has been previously validated in other mouse models of respiratory infections ^18,55,56^.

### Preparation of lung homogenates for viral plaque assay and western blot

Mouse were euthanized, the inferior vena cava was incised, and the right ventricle was perfused in situ with >1 ml of sterile PBS. Lungs were removed and kept on ice prior to and during homogenization for 30 s in PBS. The homogenate was split into two aliquots and an additional 1 mL of PBS was added to one of the aliquots, which was then centrifuged (2000 rpm for 10 min, at 4°C). MDCK cells were grown in 6-well plates to 100% confluence, then incubated with serial 10-fold dilutions of lung homogenate in DMEM and 1% bovine serum albumin (BSA) for 1 h (37°C). Supernatants were then aspirated, the cells were washed with PBS, 3 ml of replacement media [2.4% Avicel (IMCD), 2× DMEM, and 1.5 µg of N-acetyl trypsin] were added to each well, and the plates were incubated for 3 days. The overlay was then removed and viral plaques were visualized using naphthalene black dye solution (0.1% naphthalene black, 6% glacial acetic acid, 1.36% anhydrous sodium acetate) ^57^. The second aliquot of lung homogenate was mixed with 0.5 mL RIPA buffer and used for immunoblotting.

### Statistical analysis

Statistical analyses were carried out using Prism software (GraphPad Prism 9.0). Data are presented as means ± SE. Differences between two groups were assessed using a Student t test. Differences between multiple groups were assessed by ANOVA followed by the Tukey–Kramer honestly significant difference test. Levene’s test was used to analyze the homogeneity of variances. Significance was accepted at p < 0.05. Survival curves were analyzed using log-rank test.

## Authors contributions

S.M.C.M. and P.H.S.S. conceived and designed the research. S.M.C.M., F.C., F.J.G.G, A.N and H.A.V. performed the experiments. S.M.C.M. and H.M. performed the bioinformatics analysis. S.M.C.M. and P.H.S.S. analyzed and interpreted the data. G.R.S.B. and G.J.B. contributed reagents or analytic tools, S.M.C.M. and P.H.S.S. wrote the manuscript. All authors provided edits and feedback on the manuscript.

## Acknowledgments

This work was supported by R01HL131745 from the NIH and Merit Review I01 CX002350 from the Department of Veterans Affairs to P.H.S.S., and by grants U19AI35964, P01AG049665, P01AG04966506S1, and R01HL147575 from NIH and Merit Review I01 CX001777 from the Department of Veterans Affairs to G.R.S.B.

Imaging work was performed at the Northwestern University Center for Advanced Microscopy generously supported by NCI CCSG P30CA060553. Histology services were provided by the Northwestern University Mouse Histology and Phenotyping Laboratory which is supported by NCI P30-CA060553. Northwestern University Flow Cytometry Core Facility is supported by NCI Cancer Center Support Grant P01AG049665.

This research was supported in part through the computational resources and staff contributions provided by the Genomics Compute Cluster, which is jointly supported by the Feinberg School of Medicine, the Center for Genetic Medicine, and Feinberg’s Department of Biochemistry and Molecular Genetics, the Office of the Provost, the Office for Research and Northwestern Information Technology. The Genomics Compute Cluster is part of Quest, Northwestern University’s high-performance computing facility, with the purpose to advance research in genomics.

## Footnotes

1 We use *Zfhx3* and *Zfhx3* to denote the gene and protein, respectively, in both the mouse and human context. We do this for simplicity, rather than switch back and forth between the conventional designations of *Zfhx3/Zfhx3* for the mouse and *ZFHX3/ZFHX3* for the human gene and protein.

## References

1. Sethi, S., and Murphy, T.F. (2008). Infection in the pathogenesis and course of chronic obstructive pulmonary disease. N Engl J Med 359, 2355–2365. 10.1056/NEJMra0800353.

2. Mallia, P., and Johnston, S.L. (2007). Influenza infection and COPD. Int J Chron Obstruct Pulmon Dis 2, 55–64.

3. De Serres, G., Lampron, N., La Forge, J., Rouleau, I., Bourbeau, J., Weiss, K., Barret, B., and Boivin, G. (2009). Importance of viral and bacterial infections in chronic obstructive pulmonary disease exacerbations. J Clin Virol 46, 129–133. 10.1016/j.jcv.2009.07.010.

4. Gerke, A.K., Tang, F., Yang, M., Foster, E.D., Cavanaugh, J.E., and Polgreen, P.M. (2013). Predicting chronic obstructive pulmonary disease hospitalizations based on concurrent influenza activity. COPD 10, 573–580. 10.3109/15412555.2013.777400.

5. Moser, K.M., Shibel, E.M., and Beamon, A.J. (1973). Acute respiratory failure in obstructive lung disease. Long-term survival after treatment in an intensive care unit. JAMA 225, 705–707.

6. Martin, T.R., Lewis, S.W., and Albert, R.K. (1982). The prognosis of patients with chronic obstructive pulmonary disease after hospitalization for acute respiratory failure. Chest 82, 310–314.

7. Goel, A., Pinckney, R.G., and Littenberg, B. (2003). APACHE II predicts long-term survival in COPD patients admitted to a general medical ward. Journal of general internal medicine 18, 824–830.

8. Groenewegen, K.H., Schols, A.M., and Wouters, E.F. (2003). Mortality and mortality-related factors after hospitalization for acute exacerbation of COPD. Chest 124, 459–467.

9. Mohan, A., Premanand, R., Reddy, L.N., Rao, M.H., Sharma, S.K., Kamity, R., and Bollineni, S. (2006). Clinical presentation and predictors of outcome in patients with severe acute exacerbation of chronic obstructive pulmonary disease requiring admission to intensive care unit. BMC pulmonary medicine 6, 27. 10.1186/1471-2466-6-27.

10. Sin, D.D., Man, S.F., and Marrie, T.J. (2005). Arterial carbon dioxide tension on admission as a marker of in-hospital mortality in community-acquired pneumonia. Am J Med 118, 145–150. 10.1016/j.amjmed.2004.10.014.

11. Murtagh, P., Giubergia, V., Viale, D., Bauer, G., and Pena, H.G. (2009). Lower respiratory infections by adenovirus in children. Clinical features and risk factors for bronchiolitis obliterans and mortality. Pediatr Pulmonol 44, 450–456. 10.1002/ppul.20984.

12. Belkin, R.A., Henig, N.R., Singer, L.G., Chaparro, C., Rubenstein, R.C., Xie, S.X., Yee, J.Y., Kotloff, R.M., Lipson, D.A., and Bunin, G.R. (2006). Risk factors for death of patients with cystic fibrosis awaiting lung transplantation. American journal of respiratory and critical care medicine 173, 659–666. 10.1164/rccm.200410-1369OC.

13. Helenius, I.T., Krupinski, T., Turnbull, D.W., Gruenbaum, Y., Silverman, N., Johnson, E.A., Sporn, P.H., Sznajder, J.I., and Beitel, G.J. (2009). Elevated CO2 suppresses specific Drosophila innate immune responses and resistance to bacterial infection. Proc Natl Acad Sci U S A 106, 18710–18715. 10.1073/pnas.0905925106.

14. Wang, N., Gates, K.L., Trejo, H., Favoreto, S., Jr., Schleimer, R.P., Sznajder, J.I., Beitel, G.J., and Sporn, P.H. (2010). Elevated CO2 selectively inhibits interleukin-6 and tumor necrosis factor expression and decreases phagocytosis in the macrophage. FASEB J 24, 2178–2190. 10.1096/fj.09-136895.

15. Casalino-Matsuda, S.M., Berdnikovs, S., Wang, N., Nair, A., Gates, K.L., Beitel, G.J., and Sporn, P.H.S. (2021). Hypercapnia selectively modulates LPS-induced changes in innate immune and DNA replication-related gene transcription in the macrophage. Interface Focus 11, 20200039. 10.1098/rsfs.2020.0039.

16. Casalino-Matsuda, S.M., Nair, A., Beitel, G.J., Gates, K.L., and Sporn, P.H. (2015). Hypercapnia Inhibits Autophagy and Bacterial Killing in Human Macrophages by Increasing Expression of Bcl-2 and Bcl-xL. J Immunol 194, 5388–5396. 10.4049/jimmunol.1500150.

17. Gates, K.L., Howell, H.A., Nair, A., Vohwinkel, C.U., Welch, L.C., Beitel, G.J., Hauser, A.R., Sznajder, J.I., and Sporn, P.H. (2013). Hypercapnia impairs lung neutrophil function and increases mortality in murine pseudomonas pneumonia. Am J Respir Cell Mol Biol 49, 821–828. 10.1165/rcmb.2012-0487OC.

18. Casalino-Matsuda, S.M., Chen, F., Gonzalez-Gonzalez, F.J., Nair, A., Dib, S., Yemelyanov, A., Gates, K.L., Budinger, G.R.S., Beitel, G.J., and Sporn, P.H.S. (2020). Hypercapnia Suppresses Macrophage Antiviral Activity and Increases Mortality of Influenza A Infection via Akt1. J Immunol 205, 489–501. 10.4049/jimmunol.2000085.

19. Helenius, I.T., Haake, R.J., Kwon, Y.J., Hu, J.A., Krupinski, T., Casalino-Matsuda, S.M., Sporn, P.H.S., Sznajder, J.I., and Beitel, G.J. (2016). Identification of Drosophila Zfh2 as a Mediator of Hypercapnic Immune Regulation by a Genome-Wide RNA Interference Screen. J Immunol 196, 655–667. 10.4049/jimmunol.1501708.

20. Miura, Y., Tam, T., Ido, A., Morinaga, T., Miki, T., Hashimoto, T., and Tamaoki, T. (1995). Cloning and characterization of an ATBF1 isoform that expresses in a neuronal differentiation-dependent manner. J Biol Chem 270, 26840–26848. 10.1074/jbc.270.45.26840.

21. Ishii, Y., Kawaguchi, M., Takagawa, K., Oya, T., Nogami, S., Tamura, A., Miura, Y., Ido, A., Sakata, N., Hashimoto-Tamaoki, T., et al. (2003). ATBF1-A protein, but not ATBF1-B, is preferentially expressed in developing rat brain. J Comp Neurol 465, 57–71. 10.1002/cne.10807.

22. Sun, X., Frierson, H.F., Chen, C., Li, C., Ran, Q., Otto, K.B., Cantarel, B.L., Vessella, R.L., Gao, A.C., Petros, J., et al. (2005). Frequent somatic mutations of the transcription factor ATBF1 in human prostate cancer. Nat Genet 37, 407–412. 10.1038/ng1528.

23. Benjamin, E.J., Rice, K.M., Arking, D.E., Pfeufer, A., van Noord, C., Smith, A.V., Schnabel, R.B., Bis, J.C., Boerwinkle, E., Sinner, M.F., et al. (2009). Variants in ZFHX3 are associated with atrial fibrillation in individuals of European ancestry. Nat Genet 41, 879–881. 10.1038/ng.416.

24. Sun, X., Fu, X., Li, J., Xing, C., Martin, D.W., Zhang, H.H., Chen, Z., and Dong, J.T. (2012). Heterozygous deletion of Atbf1 by the Cre-loxP system in mice causes preweaning mortality. Genesis 50, 819–827. 10.1002/dvg.22041.

25. Kim, H.M., Lee, Y.W., Lee, K.J., Kim, H.S., Cho, S.W., van Rooijen, N., Guan, Y., and Seo, S.H. (2008). Alveolar macrophages are indispensable for controlling influenza viruses in lungs of pigs. Journal of virology 82, 4265–4274. 10.1128/JVI.02602-07.

26. Tate, M.D., Pickett, D.L., van Rooijen, N., Brooks, A.G., and Reading, P.C. (2010). Critical role of airway macrophages in modulating disease severity during influenza virus infection of mice. Journal of virology 84, 7569–7580. 10.1128/JVI.00291-10.

27. Purnama, C., Ng, S.L., Tetlak, P., Setiagani, Y.A., Kandasamy, M., Baalasubramanian, S., Karjalainen, K., and Ruedl, C. (2014). Transient ablation of alveolar macrophages leads to massive pathology of influenza infection without affecting cellular adaptive immunity. Eur J Immunol 44, 2003–2012. 10.1002/eji.201344359.

28. Schneider, C., Nobs, S.P., Heer, A.K., Kurrer, M., Klinke, G., van Rooijen, N., Vogel, J., and Kopf, M. (2014). Alveolar macrophages are essential for protection from respiratory failure and associated morbidity following influenza virus infection. PLoS Pathog 10, e1004053. 10.1371/journal.ppat.1004053.

29. McCubbrey, A.L., Allison, K.C., Lee-Sherick, A.B., Jakubzick, C.V., and Janssen, W.J. (2017). Promoter Specificity and Efficacy in Conditional and Inducible Transgenic Targeting of Lung Macrophages. Front Immunol 8, 1618. 10.3389/fimmu.2017.01618.

30. Fortini, M.E., Lai, Z.C., and Rubin, G.M. (1991). The Drosophila zfh-1 and zfh-2 genes encode novel proteins containing both zinc-finger and homeodomain motifs. Mech Dev 34, 113–122. 10.1016/0925-4773(91)90048-b.

31. Yasuda, H., Mizuno, A., Tamaoki, T., and Morinaga, T. (1994). ATBF1, a multiple-homeodomain zinc finger protein, selectively down-regulates AT-rich elements of the human alpha-fetoprotein gene. Mol Cell Biol 14, 1395–1401. 10.1128/mcb.14.2.1395-1401.1994.

32. Kaspar, P., Dvorakova, M., Kralova, J., Pajer, P., Kozmik, Z., and Dvorak, M. (1999). Myb-interacting protein, ATBF1, represses transcriptional activity of Myb oncoprotein. J Biol Chem 274, 14422–14428. 10.1074/jbc.274.20.14422.

33. Sun, X., Zhou, Y., Otto, K.B., Wang, M., Chen, C., Zhou, W., Subramanian, K., Vertino, P.M., and Dong, J.T. (2007). Infrequent mutation of ATBF1 in human breast cancer. J Cancer Res Clin Oncol 133, 103–105. 10.1007/s00432-006-0148-y.

34. Mabuchi, M., Kataoka, H., Miura, Y., Kim, T.S., Kawaguchi, M., Ebi, M., Tanaka, M., Mori, Y., Kubota, E., Mizushima, T., et al. (2010). Tumor suppressor, AT motif binding factor 1 (ATBF1), translocates to the nucleus with runt domain transcription factor 3 (RUNX3) in response to TGF-beta signal transduction. Biochem Biophys Res Commun 398, 321–325. 10.1016/j.bbrc.2010.06.090.

35. Duan, M., Hibbs, M.L., and Chen, W. (2017). The contributions of lung macrophage and monocyte heterogeneity to influenza pathogenesis. Immunol Cell Biol 95, 225–235. 10.1038/icb.2016.97.

36. Ji, Z.X., Wang, X.Q., and Liu, X.F. (2021). NS1: A Key Protein in the “Game” Between Influenza A Virus and Host in Innate Immunity. Front Cell Infect Microbiol 11, 670177. 10.3389/fcimb.2021.670177.

37. Sountoulidis, A., Marco Salas, S., Braun, E., Avenel, C., Bergenstrahle, J., Theelke, J., Vicari, M., Czarnewski, P., Liontos, A., Abalo, X., et al. (2023). A topographic atlas defines developmental origins of cell heterogeneity in the human embryonic lung. Nat Cell Biol 25, 351–365. 10.1038/s41556-022-01064-x.

38. Tabula Muris, C., Overall, c., Logistical, c., Organ, c., processing, Library, p., sequencing, Computational data, a., Cell type, a., Writing, g., et al. (2018). Single-cell transcriptomics of 20 mouse organs creates a Tabula Muris. Nature 562, 367–372. 10.1038/s41586-018-0590-4.

39. Travaglini, K.J., Nabhan, A.N., Penland, L., Sinha, R., Gillich, A., Sit, R.V., Chang, S., Conley, S.D., Mori, Y., Seita, J., et al. (2020). A molecular cell atlas of the human lung from single-cell RNA sequencing. Nature 587, 619–625. 10.1038/s41586-020-2922-4.

40. Casalino-Matsuda, S.M., Wang, N., Ruhoff, P.T., Matsuda, H., Nlend, M.C., Nair, A., Szleifer, I., Beitel, G.J., Sznajder, J.I., and Sporn, P.H.S. (2018). Hypercapnia Alters Expression of Immune Response, Nucleosome Assembly and Lipid Metabolism Genes in Differentiated Human Bronchial Epithelial Cells. Scientific Reports 8, 13508. 10.1038/s41598-018-32008-x.

41. King, D.T., Zhu, S., Hardie, D.B., Serrano-Negron, J.E., Madden, Z., Kolappan, S., and Vocadlo, D.J. (2022). Chemoproteomic identification of CO(2)-dependent lysine carboxylation in proteins. Nat Chem Biol 18, 782–791. 10.1038/s41589-022-01043-1.

42. Linthwaite, V.L., Janus, J.M., Brown, A.P., Wong-Pascua, D., O’Donoghue, A.C., Porter, A., Treumann, A., Hodgson, D.R.W., and Cann, M.J. (2018). The identification of carbon dioxide mediated protein post-translational modifications. Nat Commun 9, 3092. 10.1038/s41467-018-05475-z.

43. Tomlinson, G.S., Booth, H., Petit, S.J., Potton, E., Towers, G.J., Miller, R.F., Chain, B.M., and Noursadeghi, M. (2012). Adherent human alveolar macrophages exhibit a transient pro-inflammatory profile that confounds responses to innate immune stimulation. PLoS One 7, e40348. 10.1371/journal.pone.0040348.

44. Trouplin, V., Boucherit, N., Gorvel, L., Conti, F., Mottola, G., and Ghigo, E. (2013). Bone marrow-derived macrophage production. J Vis Exp, e50966. 10.3791/50966.

45. Davies, J.Q., and Gordon, S. (2005). Isolation and culture of human macrophages. Methods in molecular biology 290, 105–116.

46. Casalino-Matsuda, S.M., Forteza, R.M., and Monzon, M.E. (2008). Hyaluronan Fragments Induce MUC5B Expression through a Monocyte Chemoattractant Protein-1/ CCR2 Dependent Mechanism. Am. J. Respir. Crit. Care Med 177, A994.

47. Misharin, A.V., Morales-Nebreda, L., Mutlu, G.M., Budinger, G.R., and Perlman, H. (2013). Flow cytometric analysis of macrophages and dendritic cell subsets in the mouse lung. Am J Respir Cell Mol Biol 49, 503–510. 10.1165/rcmb.2013-0086MA.

48. Lawrence, M., Huber, W., Pages, H., Aboyoun, P., Carlson, M., Gentleman, R., Morgan, M.T., and Carey, V.J. (2013). Software for computing and annotating genomic ranges. PLoS Comput Biol 9, e1003118. 10.1371/journal.pcbi.1003118.

49. Robinson, M.D., McCarthy, D.J., and Smyth, G.K. (2010). edgeR: a Bioconductor package for differential expression analysis of digital gene expression data. Bioinformatics 26, 139–140. 10.1093/bioinformatics/btp616.

50. McCarthy, D.J., Chen, Y., and Smyth, G.K. (2012). Differential expression analysis of multifactor RNA-Seq experiments with respect to biological variation. Nucleic Acids Res 40, 4288–4297. 10.1093/nar/gks042.

51. Breuer, K., Foroushani, A.K., Laird, M.R., Chen, C., Sribnaia, A., Lo, R., Winsor, G.L., Hancock, R.E., Brinkman, F.S., and Lynn, D.J. (2013). InnateDB: systems biology of innate immunity and beyond--recent updates and continuing curation. Nucleic Acids Res 41, D1228–1233. 10.1093/nar/gks1147.

52. Warde-Farley, D., Donaldson, S.L., Comes, O., Zuberi, K., Badrawi, R., Chao, P., Franz, M., Grouios, C., Kazi, F., Lopes, C.T., et al. (2010). The GeneMANIA prediction server: biological network integration for gene prioritization and predicting gene function. Nucleic Acids Res 38, W214–220. 10.1093/nar/gkq537.

53. Shannon, P., Markiel, A., Ozier, O., Baliga, N.S., Wang, J.T., Ramage, D., Amin, N., Schwikowski, B., and Ideker, T. (2003). Cytoscape: a software environment for integrated models of biomolecular interaction networks. Genome Res 13, 2498–2504. 10.1101/gr.1239303.

54. Cimolai, N., Taylor, G.P., Mah, D., and Morrison, B.J. (1992). Definition and application of a histopathological scoring scheme for an animal model of acute Mycoplasma pneumoniae pulmonary infection. Microbiol Immunol 36, 465–478.

55. Alsuwaidi, A.R., George, J.A., Almarzooqi, S., Hartwig, S.M., Varga, S.M., and Souid, A.K. (2017). Sirolimus alters lung pathology and viral load following influenza A virus infection. Respir Res 18, 136. 10.1186/s12931-017-0618-6.

56. Martin, R.J., Chu, H.W., Honour, J.M., and Harbeck, R.J. (2001). Airway inflammation and bronchial hyperresponsiveness after Mycoplasma pneumoniae infection in a murine model. Am J Respir Cell Mol Biol 24, 577–582. 10.1165/ajrcmb.24.5.4315.

57. Jing, X., Ma, C., Ohigashi, Y., Oliveira, F.A., Jardetzky, T.S., Pinto, L.H., and Lamb, R.A. (2008). Functional studies indicate amantadine binds to the pore of the influenza A virus M2 proton-selective ion channel. Proc Natl Acad Sci U S A 105, 10967–10972. 10.1073/pnas.0804958105.

