## Supplementary material for "Myeloid *Zfhx3* Deficiency Protects Against Hypercapnia-induced Suppression of Host Defense Against Influenza A Virus"

**Supplementary Materials:**

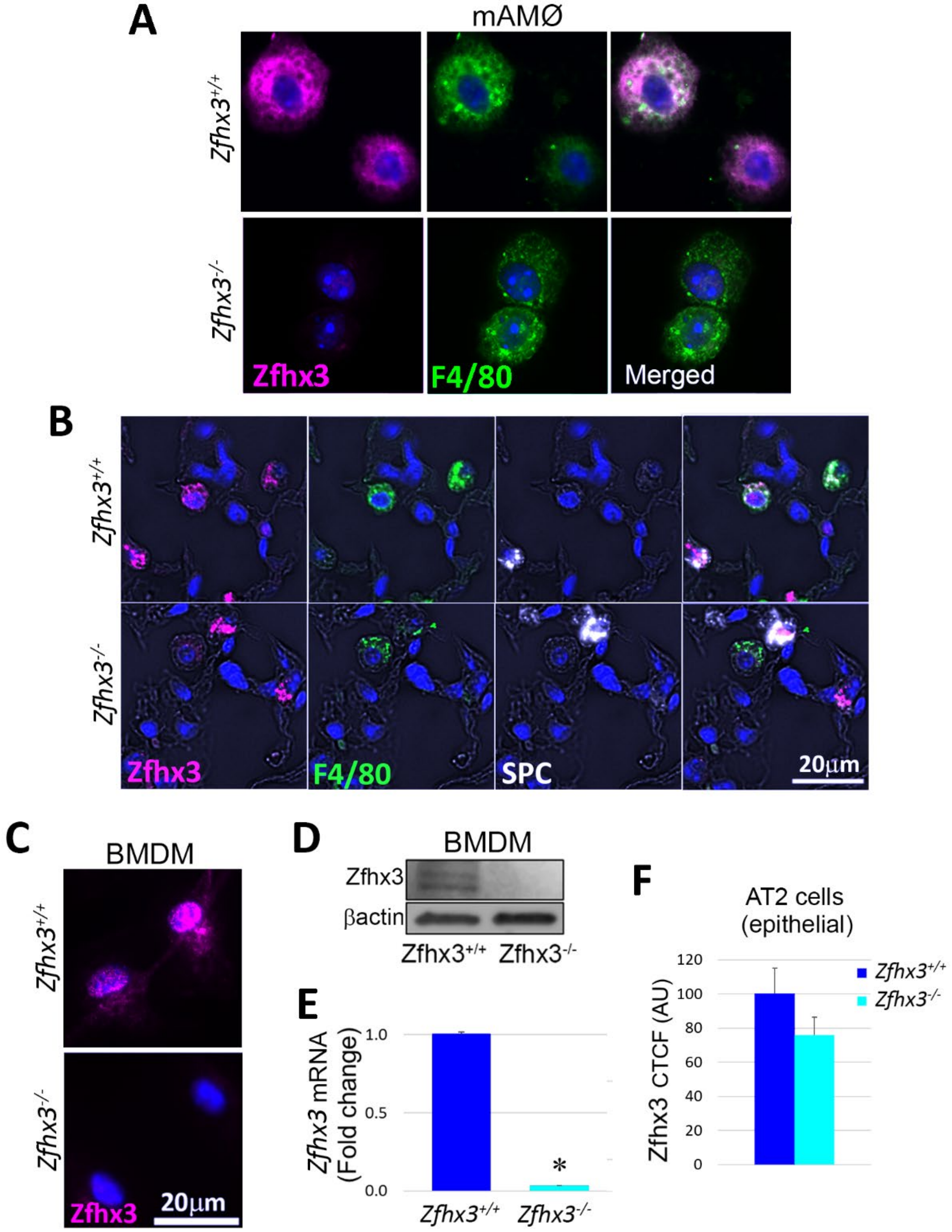


**Supplemental Figure 1:** ***Zfhx3* protein and mRNA are markedly reduced in alveolar macrophages from *Zfhx3*$^{fl/fl}$*LyzM*$^{Cre}$ (*Zfhx3*$^{-/-}$) mice.** BAL AMØs, BMDM and lungs were obtained from *Zfhx3*$^{+/+}$ and *Zfhx3*$^{-/-}$ mice. AMΦs (A), lung tissue sections (B) and BMDM (C) stained for Zfhx3 (magenta), F4/80 (green, MØs) and SPC (white, alveolar epithelial cells). Nuclei stained with

DAPI (blue). Zfhx3 protein (D) and *Zfhx3* mRNA (E) expression in BMDMs measured by immunoblot and qPCR respectively. *P<0.001 vs *Zfhx3*$^{+/+}$. Zfhx3 expression by immunofluorescence and measured as corrected total cell fluorescence (CTCF, expressed in arbitrary units (AU)) in AT2 cells (F).

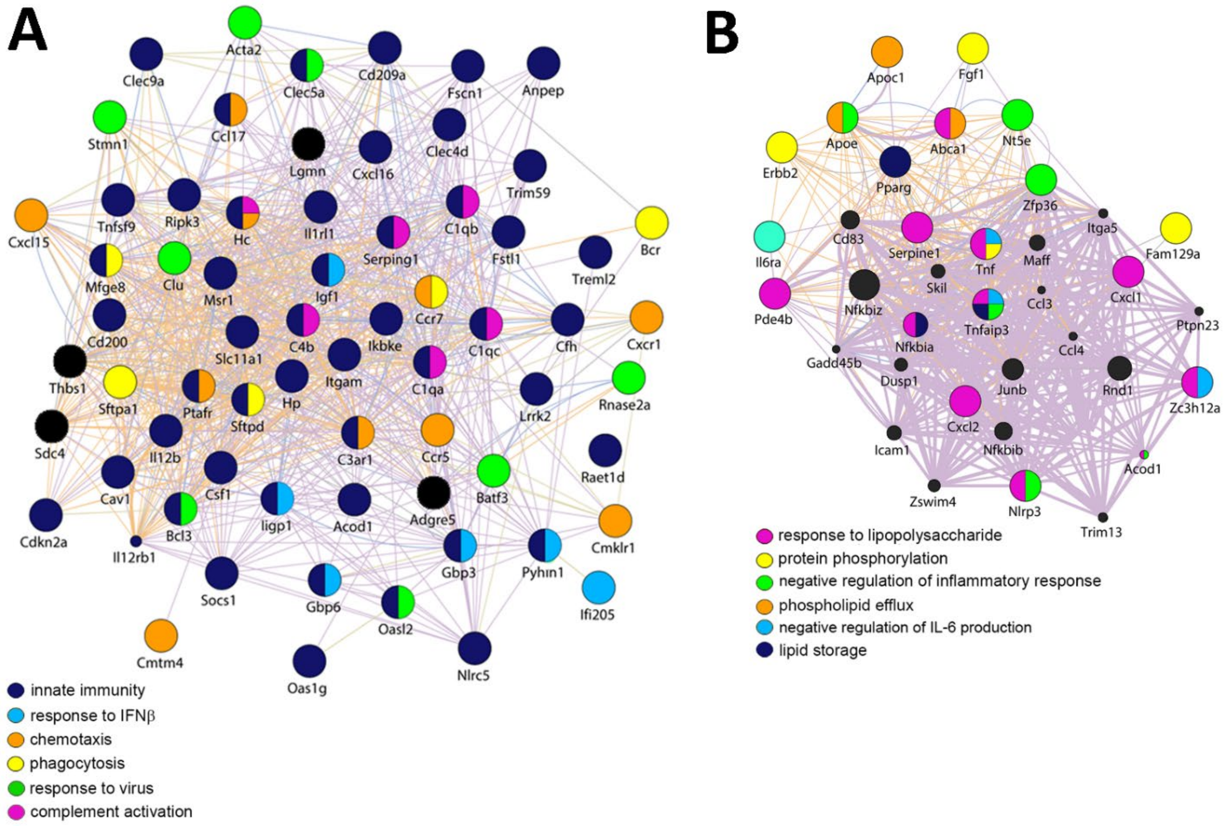


<u>Supplemental Figure 2</u>: **Gene networks downregulated and upregulated in alveolar macrophages from mice exposed to 10% $CO_2$ for 7 days.** RNA-seq analysis was performed in AMØs obtained from mice exposed to room air or 10% $CO_2$ (hypercapnia, HC) for 7 days and isolated by flow cytometry. Gene networks of GO biological processes downregulated (A) or upregulated (B) by hypercapnia.

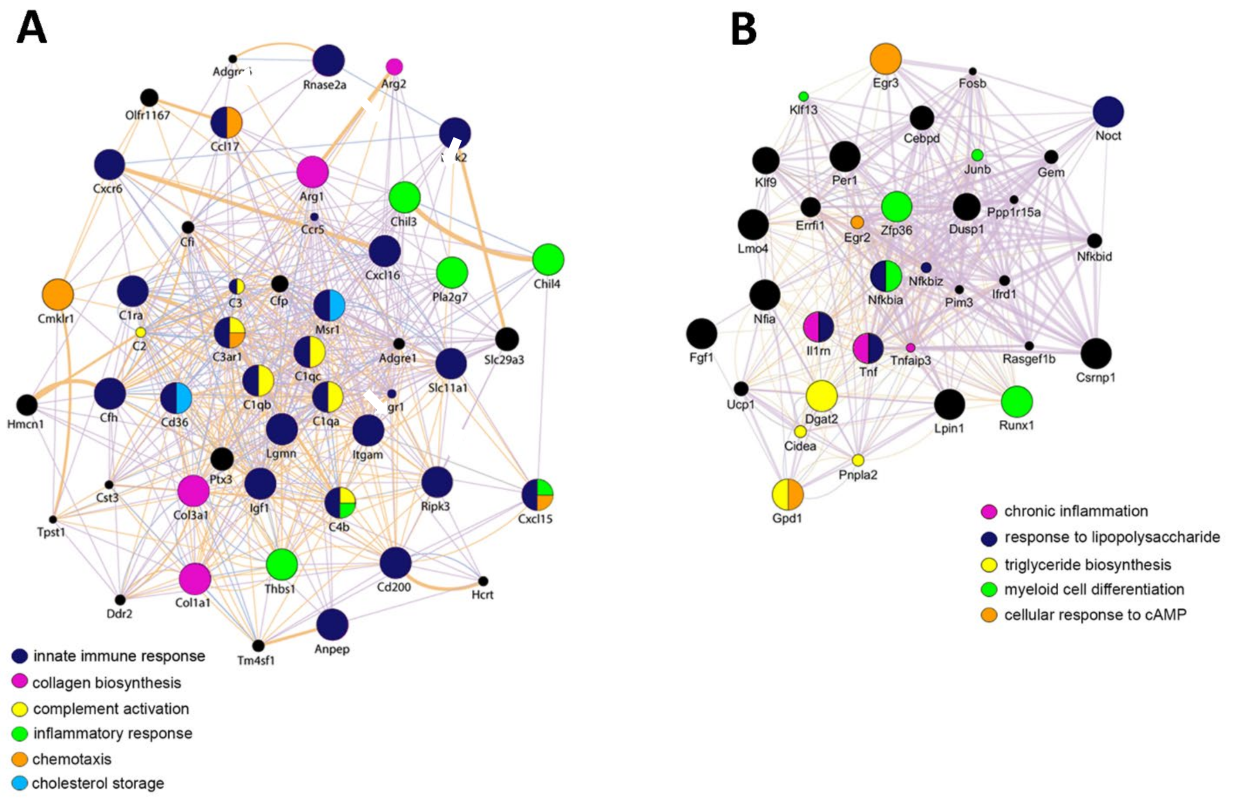


**Supplemental Figure 3: Gene networks for which myeloid *Zfhx3* deficiency abrogates hypercapnia-induced transcriptional alterations in alveolar macrophages.** *Zfhx3*$^{+/+}$ and *Zfhx3*$^{-/-}$ mice were exposed to room air or 10% $CO_2$ (hypercapnia, HC) for 7 days. AMØs from those mice were isolated by flow cytometry assessed by RNAseq analysis. Gene networks of GO biological processes in cluster 1 (A) and cluster 2 (B).

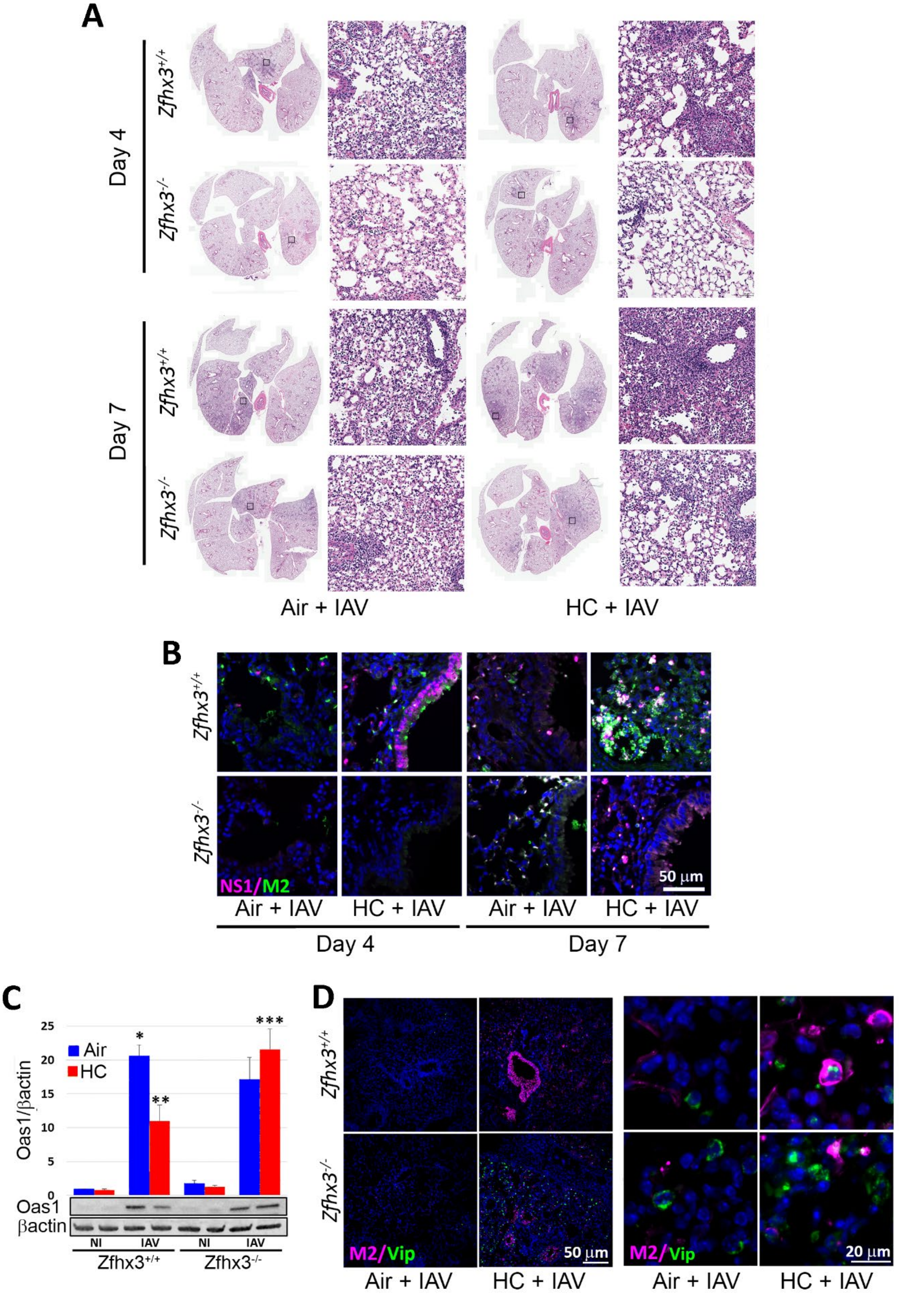
A
Day 4
Day 7
Zfhx3+/+
Zfhx3-/-
Air + IAV
HC + IAV
B
NS1/M2
50 μm
Day 4
Day 7
C
Oas1/βactin
Air
HC
25
20
15
10
5
0
*
**
***
Oas1
βactin
NI
IAV
Zfhx3+/+
Zfhx3-/-
D
M2/Vip
50 μm
20 μm

**<u>Supplemental Figure 4</u>: Myeloid *Zfhx3* deficiency protects against hypercapnia-induced suppression of antiviral gene and protein expression and increases in viral protein expression and lung injury in IAV-infected mice (3 pfu/mouse).** *Zfhx3*$^{+/+}$ and *Zfhx3*$^{-/-}$ mice pre-exposed to normoxic hypercapnia (10% $CO_2$/21% $O_2$, HC) for 3 days or air as control, then infected intratracheally with 3 pfu IAV (A/WSN/33) per animal. Lungs from IAV-infected mice harvested 4 and 7 dpi were sectioned and stained with H&E (A). Expression of viral NS1 (magenta) and M2 (green) protein was assessed in lung tissue sections from mice sacrificed 4 and 7 dpi (B). Oas1 protein expression from homogenized lung tissue at 4 dpi assessed by immunoblot (C). Expression of viperin (green) transcripts and viral M2 (magenta) protein were assessed by RNAscope and immunofluorescence respectively in lung tissue sections from mice sacrificed 4 dpi (D). *P<0.001 vs *Zfhx3*$^{+/+}$ Air; **p<0.05 vs *Zfhx3*$^{+/+}$ Air + IAV, and **p<0.05 vs *Zfhx3*$^{+/+}$ HC + IAV. Nuclei were stained with DAPI (blue).

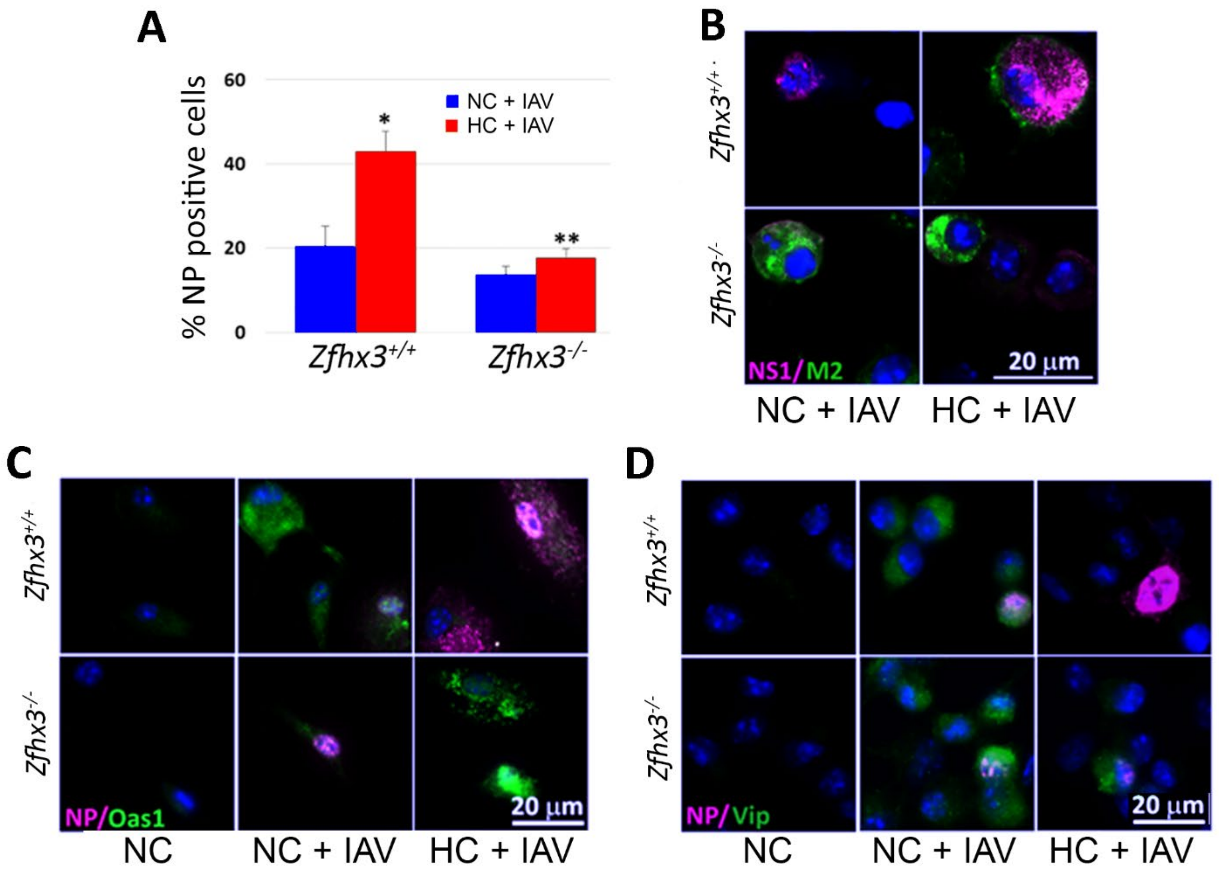


**<u>Supplemental Figure 5:</u> Myeloid *Zfhx3* deficiency increased viral replication caused by hypercapnia and abrogates suppression of antiviral protein expression in IAV-infected mouse BMDM.** *Zfhx3*$^{+/+}$ and *Zfhx3*$^{-/-}$ BMDM cultured under (5% $CO_2$/95% air, NC) or hypercapnic (15% $CO_2$/21% $O_2$/64% $N_2$, HC) conditions for 18 h, infected with IAV, placed in NC or HC for additional 24 h. NP positive BMDM assessed by immunofluorescence and quantified using ImageJ (A). BMDM stained for viral NS1 (magenta) and M2 (green) (B) or stained for NP (magenta), OAS1 (green) and viperin (green) (C, D). *P<0.001 vs *Zfhx3*$^{+/+}$ NC + IAV; **p<0.05 vs *Zfhx3*$^{+/+}$ HC + IAV.

**Supplemental Table 1**

| **Antibodies** | | |
|---|---|---|
| Rabbit polyclonal anti-NS1 | GeneTex | Cat#GTX125990; RRID:AB_11170327 |
| Mouse monoclonal anti-IAV M2 | Invitrogen | Cat#MA1-082; RRID:AB_10892134 |
| Mouse monoclonal anti-IAV NP | Abcam | Cat#ab128193; RRID:AB_11143769 |
| Rat monoclonal anti-F4/80 | Abcam | Cat#ab6640; RRID:AB_1140040 |
| Goat polyclonal anti-SPC | Santa Cruz Biotechnology | Cat#Sc-7706; RRID:AB_2185507 |
| Goat polyclonal anti-Chitinase 3-like 3/ECF-L | Novus | Cat#AF2446;<br>RRID: AB_2079008 |
| Rabbit polyclonal anti-ATBF1 (D1-120) (Human) | MBL INTERNATIONAL CORP | Cat#PD010;<br>RRID: AB_10598330 |
| Rabbit polyclonal anti-OAS1 | Abcam | Cat#ab86343; RRID:AB_2158286 |
| Rabbit polyclonal anti-viperin | Abcam | Cat#ab73864; RRID:AB_1640850 |
| Goat anti-Rabbit IgG (H+L) Highly Cross-Adsorbed Secondary Antibody, Alexa Fluor Plus 555 | ThermoFisher | Cat#A32732; RRID:AB_2633281 |
| Donkey Anti-Mouse IgG (H+L) Antibody, Alexa Fluor 488 Conjugated | ThermoFisher | Cat#A21202; RRID:AB_141607 |
| Rabbit anti-Goat IgG (H+L) Superclonal(TM) Secondary Antibody, Alexa Fluor 647 | ThermoFisher | Cat#A27018; RRID:AB_2536082 |
| LI-COR IRDye 680RD Goat anti-Mouse | ThermoFisher | Cat#NC0252290 |
| Anti-rabbit IgG, HRP-linked Antibody | Cell Signaling Technology | Cat#7074; RRID:AB_2099233 |
| Anti-mouse IgG, HRP-linked Antibody | Cell Signaling Technology | Cat#7076; RRID:AB_330924 |
| Rabbit polyclonal anti-NS1 | GeneTex | Cat#GTX125990; RRID:AB_11170327 |
